# Somatic mutations and single-nucleus transcriptomics reveal uniquely human properties of neuronal aging

**DOI:** 10.64898/2026.09.22.753624

**Authors:** Emre Caglayan, Ishaan Lamba, Monica Manam Devi, Benjamin Finander, Lovelace J. Luquette, David Exposito-Alonso, Sijing Zhao, Bowen Jin, Michael B. Miller, Peter J. Park, Chet C. Sherwood, Christopher A. Walsh

**Affiliations:** Division of Genetics and Genomics, Manton Center for Orphan Disease Research, Boston Children’s Hospital, Boston, MA, USA; Department of Pediatrics, Harvard Medical School, Boston, MA, USA; Broad Institute of MIT and Harvard, Cambridge, MA, USA; Boston Children’s Hospital, Howard Hughes Medical Institute, Boston, MA, USA; Department of Biomedical Informatics, Harvard Medical School, Boston, MA, USA; Division of Neuropathology, Department of Pathology, Brigham and Women’s Hospital, Harvard Medical School, Boston; Department of Anthropology and Center for the Advanced Study of Human Paleobiology, The George Washington University, Washington, DC, USA

## Abstract

Human brain aging is frequently studied in model mammals that rarely show the natural age-related neurodegenerative conditions that afflict humans, yet little is known about how genomic stability compares across species during aging. Although somatic single nucleotide variant (SNV) and short insertion/deletion (Indel) mutations accumulate at rates that inversely scale to lifespan in colon cells—leaving diverse mammals with similar end-of-life burdens of mutations -- here we show that cerebral cortical neurons accumulate SNVs at annual rates that are remarkably similar across six mammalian species (human, chimpanzee, rhesus macaque, marmoset, ferret, mouse) resulting in >12-fold more mutations in human neurons at the end of life compared to mouse neurons. Despite the conservation of overall annual mutation accumulation rates, mutational patterns--and hence likely mutagenic mechanisms--show sharp differences between species, with a nucleotide substitution pattern linked to neurodegenerative diseases accumulating almost exclusively in humans during aging. Coinciding with their higher mutational burden, single-nucleus transcriptomic analyses reveal pervasive age-associated proteostasis and mitochondrial dysregulation in human neurons that is far less pronounced in aged chimpanzees and rhesus macaques. These findings show that the pervasive effects of human neuronal aging might be linked to our long lifespan and are therefore not evident in animal models.

## Introduction

Despite the efficiency of DNA repair, DNA damage leads to irreversible somatic mutations that accumulate in all cells of our body throughout life^1,2^, even in cerebral cortical neurons that are permanently postmitotic^1^. Given that species lifespan can vary up to 100-fold in mammals^3^, long-lived species could be under negative selection for somatic mutation rates, restricting their genomic contribution to age-related cellular decline. Indeed, a recent study of intestinal crypts showed that mammalian species with longer lifespans accumulate somatic mutations at a slower rate such that the end of lifespan mutation burden varies only modestly among species with >30-fold differences in lifespan, therefore suggesting a potential evolutionary “ceiling” on somatic mutation levels^4^. Other studies have uncovered evolutionary novelties in single genes that boost the efficiency of DNA repair in species with long lifespan^5–7^, suggesting that annual rates of accumulation of somatic mutations may play at least some role in life span.

Humans have the potential to live decades longer than our closest relatives, such as chimpanzees and other great apes, and accordingly might have been subject to evolutionary pressure to lower age-related mutation accumulation in our DNA. Retaining a faithful copy of DNA throughout life could be particularly important in neurons of the brain given that most neurons are not replenished during postnatal life, so that unrepaired mutations inexorably accumulate without the opportunity to replace damaged neurons from undamaged stem cells. Given the importance of DNA quality for neuronal function^8^ and the elevated susceptibility of humans to neurodegenerative disease during aging^9–12^, a systematic comparison of genomic changes in the neurons from humans and other species is critical.

To address this, we performed single-cell whole genome sequencing (scWGS) of neurons in a phylogenetically diverse sample including human, chimpanzee, rhesus macaque, marmoset, mouse and ferret to compare their somatic mutational landscapes during aging. We find that overall annual rates of somatic mutational accumulation are remarkably similar across species in two broad classifications of neuronal cell types (glutamatergic and GABAergic neurons), resulting in vastly differing end-of-lifespan mutational burdens among species. Comparative single-nucleus RNA-sequencing further suggests that human aging results in far greater levels of transcriptomic changes than its close relatives, creating a uniquely human late-life condition of widespread somatic mutation and transcriptional alteration.

## Results

### A new method to accurately identify somatic mutations across species

To identify age-related somatic mutations in single-cell whole genome data across a wide range of species, we modified an existing algorithm developed for use in human cells only, SCAN2^13^, to create a new informatic pipeline that can be applied to any species, called SCAN2-Zoo. To accurately call somatic mutations from scWGS, SCAN2 builds allelic balance curves from heterozygous SNPs (hetSNPs) to test whether candidate somatic mutations follow a similar distribution to hetSNPs in either of the phased alleles^13,14^. While hetSNPs are abundant in most species, allelic phasing is challenging due to the scarcity of population genetics datasets, preventing the algorithm’s usage for other species. To overcome this limitation in SCAN2-Zoo, we reasoned that phasing can be replaced with maximum variant allele frequency (maxVAF) at the hetSNP sites to build an AB curve, similar to a previous algorithm^15^, without compromising the accuracy of somatic mutation calls (see Methods and Supplementary Figure 1a-b for details).

We compared SCAN2-Zoo with the previously published SCAN2 results on 52 neurons across 17 neurotypical human controls^13^ to evaluate its concordance with previous results. Genome-wide mutation burdens and mutation rates per year were remarkably similar between the SCAN2-Zoo and SCAN2 for both SNVs (single nucleotide variants) and indels (insertions and deletions) **(Supplementary Figure 1c-d).** About 80% of mutation calls from either tool were exactly recapitulated in the other **(Supplementary Figure 1e-f)**. We reasoned that the remaining ∼20% calls are not identical likely due to SCAN2’s stringent criteria for mutation calling that results in false negatives that fail the stringent filters by a small margin. Indeed, the majority of such calls displayed p-values just below the threshold **(Supplementary Figure 1g)**. To further evaluate these non-identical mutations, we built mutation profiles of both sets of non-identical mutation calls as well as the identical mutation calls that revealed very high similarity between all sets **(Supplementary Figure 2)**. Together, these results indicate that SCAN2-Zoo recapitulates SCAN2 results with high fidelity while being compatible with nonhuman diploid species **(Supplementary Table 1)**.

### Uncovering mutational profiles in diverse neuronal subtypes

Because neurons are highly heterogeneous, cross-species comparisons may be intrinsically confounded by the variability across distinct cell types. To minimize this, and to explore cell type specific mutational patterns, we developed a sorting strategy to isolate glutamatergic and GABAergic neurons from human, chimpanzee, rhesus macaque and marmoset cortex by staining cortical nuclei with antibodies against NeuN and SOX6 proteins (**Figure 1a**). We validated our sorting strategy by performing single-nucleus RNA sequencing (snRNA-seq) on the sorted populations that revealed >95% purity of glutamatergic or GABAergic neurons in all species (**Figure 1b-d, Supplementary Figure 3-4)**. As expected from *SOX6* expression, GABAergic neurons were largely parvalbumin and somatostatin expressing neurons that are derived from the medial ganglionic eminence (MGE), hereafter referred to as MGE-GABAergic neurons **(Supplementary Figure 4)**.

**Figure 1:**
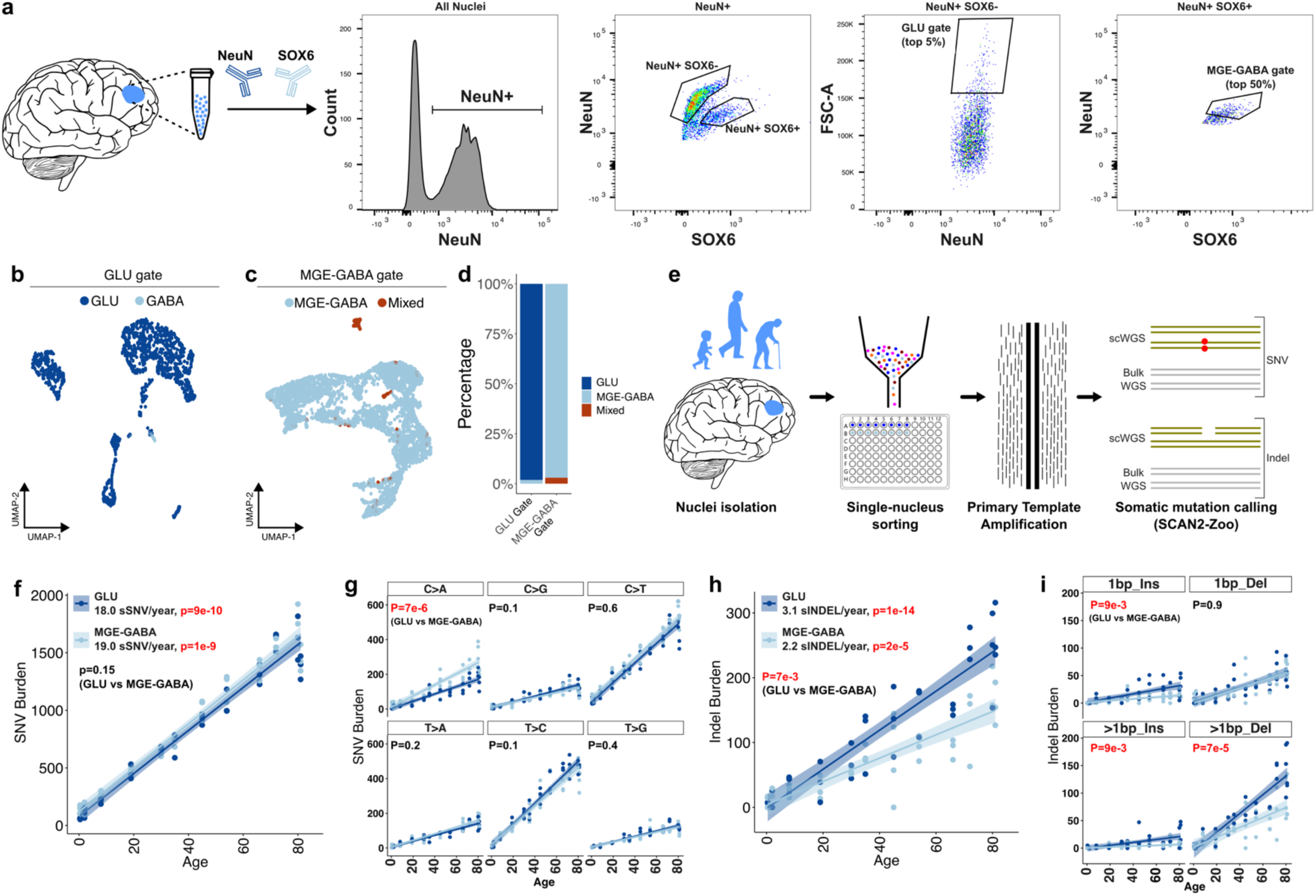
Distinct cerebral cortical neuronal subtypes display subtle yet specific differences in their mutational landscape. (a) Schematics of nuclei isolation, antibody staining and fluorescence activated nuclei sorting strategy. NeuN and SOX6 indicate antibodies against NeuN and SOX6 proteins. FSC-A is forward scatter that indicates the size of events (plots were generated using FlowJo version 10). (b-c) Single-nucleus RNA-sequencing results of nuclei sorted from (b) GLU and (c) MGE-GABA gates in panel a. (d) Percentage breakdown of cell type annotations in both GLU and MGE-GABA gates. (e) Schematics of experimental plan to perform single-cell whole genome sequencing of glutamatergic and MGE-GABAergic cells in human dorsolateral prefrontal cortex across ages. (f) Genomewide SNV burden estimates of PTA amplified glutamatergic and MGE-GABAergic cells across ages. SNV rate per year was calculated as the coefficient in linear mixed model (burden ∼ Age + (1|Subject)) for each cell type. Shaded area shows 95% confidence interval. P-value at the bottom shows the p-value for cell-type specificity of age-related change in mutation burden (burden ∼ Age*CellType + (1|Subject)). (g-i) Same as panel f, but for (g) per SNV mutational spectrum, (h) overall indel rates or (i) per indel mutational spectrum. Significant p-values (P<0.05) are shown in red.

We performed single-nucleus sorting of 37 glutamatergic cells and 39 MGE-GABAergic cells across 12 neurotypical human donors from dorsolateral prefrontal cortex, followed by single-cell whole genome amplification (scWGA) with primary template amplification (PTA). We then sequenced each scWGA PTA library at 25-30X coverage and called somatic mutations with SCAN2-Zoo (**Figure 1e, Supplementary Table 2)**. Our results revealed that mutation burden increased linearly with age at 18 SNVs/year and 3.1 indels/year in glutamatergic neurons (**Figure 1f,g**), similar to previous results^2,13^. MGE-GABAergic neurons accumulated SNVs at a slightly faster rate than glutamatergic neurons, although this difference was not significant at the sample size studied (p=0.15) (**Figure 1f**). Intriguingly, this trend was mostly driven by C>A transversions that accumulated 34% faster (p=7e-6) in MGE-GABAergic neurons than in glutamatergic neurons (**Figure 1g**). In contrast to SNVs, indels accumulated significantly faster in glutamatergic neurons than in MGE-GABAergic neurons (p=7e-3), mainly driven by 2bp deletions (**Figure 1h,i**). Taken together, these results reveal subtle yet specific differences in the mutational landscape of glutamatergic and MGE-GABAergic neurons in human cortex.

### Similar neuronal mutation rates in mammalian species with vastly different lifespans

Generating tissue and cell type matched scWGS data from the dorsolateral prefrontal cortex on nonhuman primates (chimpanzee: *Pan troglodytes*, rhesus macaque: *Macaca mulatta* and marmoset: *Callithrix jacchus*) across ages allowed species comparisons of mutation burdens and patterns. As expected, both SNV and indel mutation burden increased linearly with age in all species (**Figure 2, Supplementary Table 2)**. However, unlike in intestinal crypts^4^, annual rates of somatic mutation were surprisingly similar across species (**Figure 2a-h**). To test this in additional mammals, we collected scWGS and matched bulk WGS across ages in the F1 hybrids of BALB/c and C57BL/6 mouse strains (*Mus musculus*) and ferret (*Mustela putorius furo*) frontal cortex neurons. We used the same gating strategy to isolate pure glutamatergic neurons in both species **(Supplemental Figure 5)**. Mouse neurons had very low mutation burdens even in the oldest individuals (2.7 years old), with mutation burdens increasing linearly from ages 0.4 year to 2.7 years at ∼18.6 SNV/year (SEM = 5.9, p=0.03) (**Figure 2i,j**), an annual rate nominally similar to that observed in humans. We observed a similar trend in ferret glutamatergic neurons (24 SNV/year, SEM = 9.6, p = 0.02) (**Figure 2k,l**). We did not discover any indels in mouse or ferret neurons, in stark contrast with aged human neurons (which show 250-300 indels per genome^13^) but these low burdens resemble 0-2 year old human neurons that also rarely show indels. Extrapolation of end-of-lifespan mutation burden in all species showed that aging human neurons harbor hundreds more SNVs and indels than other species, even our closest relative, the chimpanzees (**Figure 2m,n, Methods)**. These results reflect remarkable species conservation of the annual rate of SNV and indel accumulation, but result in a dramatic lack of constraint on the overall end-of-life neuronal mutation burdens in humans compared to other species.

**Figure 2:**
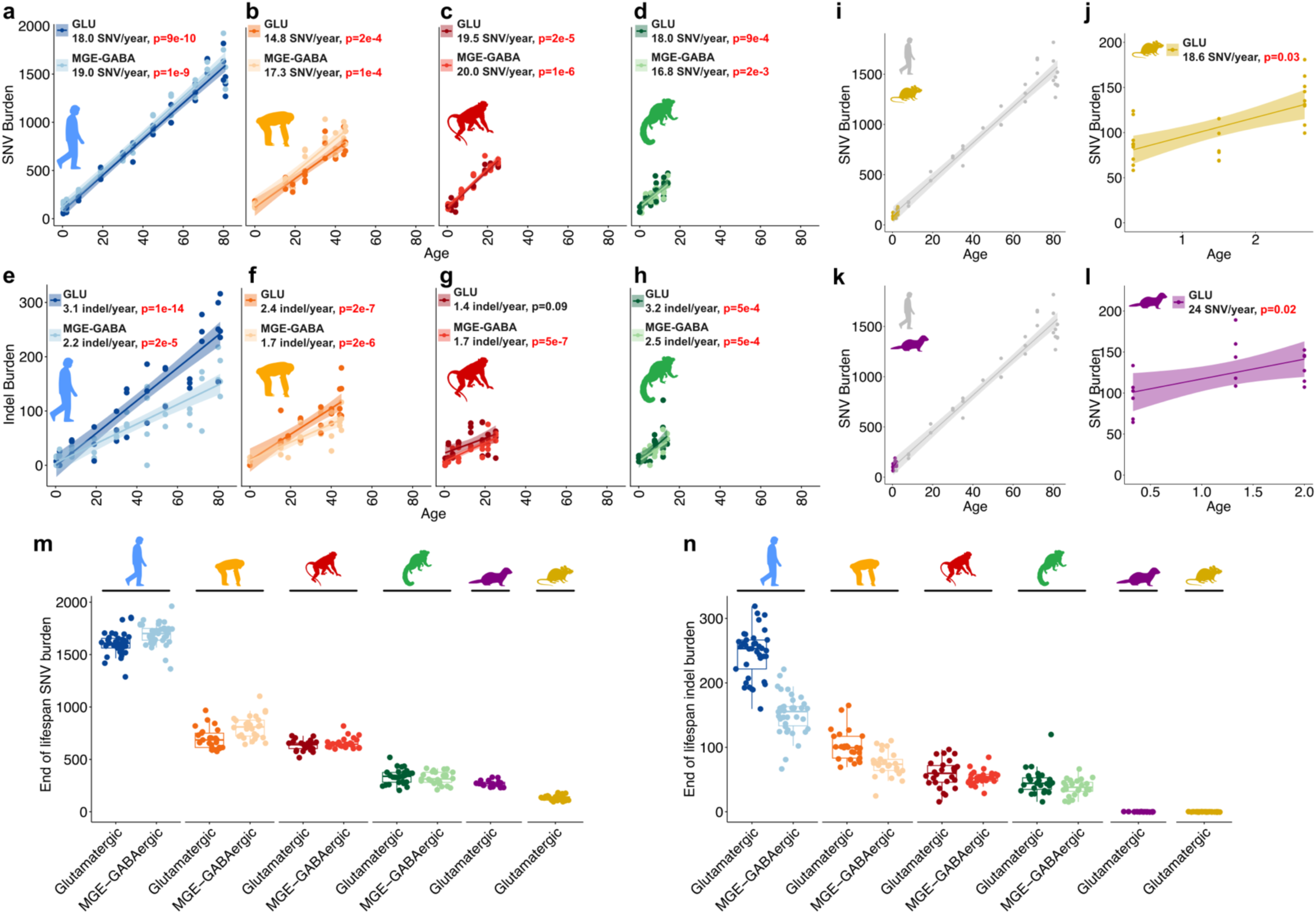
Mammalian species with vastly different lifespans accumulate somatic mutations at similar annual rates in neurons. (a-d) Genomewide mutation burden estimates for SNVs in (a) humans (37 glutamatergic, 39 MGE-GABAergic neurons), (b) chimpanzees (28 glutamatergic, 26 MGE-GABAergic neurons across 8 subjects), (c) rhesus macaques (29 glutamatergic, 26 MGE-GABAergic neurons across 9 subjects) and (d) marmosets (28 glutamatergic, 25 MGE-GABAergic neurons across 9 subjects). (e-h) Same as panels a-d but for indels. (i) Overlay of genomewide SNV burdens in human and mouse (27 glutamatergic neurons across 8 subjects). (j) Expanded view of genomewide SNV burdens in mouse glutamatergic neurons. (k-l) Same as in panels i-j but for ferret glutamatergic neurons (19 glutamatergic neurons across 6 subjects). Statistics in panels a-h,j,l were calculated as described in the legend of Figure 1. (m-n) End of lifespan mutation burden estimates for each cell across species and cell types in (m) SNVs and (n) indels. P-values test whether the mutation increase with age is significant (P<0.05 are shown in red).

The low SNV burdens in mouse neurons are consistent with a previous study that identified neuronal somatic mutations by using nuclear transfer and clonal expansion of mitral and tuft cells from olfactory bulb neurons of mice aged 4 months^16^ **(Supplemental Figure 6a**). Mutational profiles in both datasets also reveal high cosine similarity (0.87), marked by relatively greater C>T transitions than other mutations **(Supplementary Figure 6b,c)**. Therefore, our scWGS based mutational landscape of mouse neurons are within the expected range of this second method.

We further tested mutation rates between humans and nonhuman primates using a complementary “duplex” technology, multiplexed end-tagging amplification of complementary strands (META-CS), which uses the Tn5 transposase to fragment and tag the original DNA template and calls somatic mutations based on strand consensus of variants on the Watson and Crick strands^17^, unlike PTA that requires even amplification of unfragmented single-cell genomes^18^. META-CS SNVs significantly increased with age in human and rhesus macaque neurons with similar mutation rates that also resulted in higher end-of-lifespan burden estimates in human neurons **(Supplementary Figure 7, Supplementary Table 3)**. These results provide confirmation with an orthologous method that annual rates of neuronal somatic mutation accumulation are highly similar between species with widely divergent lifespans, resulting in wide variation in end-of-life mutational burden between species.

### Similarities and differences in patterns of mutational signatures across species

Despite overall similarities in annual rates of mutation accumulation, analysis of nucleotide mutational spectra across species revealed a significant increase of C>T mutation rate and a significant decrease of T>A mutation rate in humans compared to nonhuman primates, while other substitution patterns were more similar (**Figure 3a, Supplementary Table 4)**. Mutational “signature” analysis has been highly successful in identifying specific mechanisms of mutation in cancer^19^, and so we performed *de novo* signature extraction on the trinucleotide spectra of all primate species. Signature analysis revealed four distinct signatures (SBS-A, B, C and D) (**Figure 3b, Supplementary Table 4)**. While SBS-B and SBS-D were largely conserved between species, SBS-A and SBS-C accumulated significantly slower and faster in humans, respectively (**Figure 3c-f**). The overall accumulation rate of SBS-C was near zero in nonhuman primates, but it made up ∼15% of the overall mutation rate in humans. Since SBS-C is largely composed of C>T transitions, and single-cell DNA sequencing artifacts are also C>T transition at non-CpG sites^13,20^, we divided SBS-C into C>T at CpG sites and C>T at non-CpG sites. C>T at CpG sites show a similar human-specific increase to C>T at non-CpG sites, indicating that SBS-C is unlikely to be explained by any potential artifact that may be restricted to humans **(Supplementary Figure 8)**.

**Figure 3:**
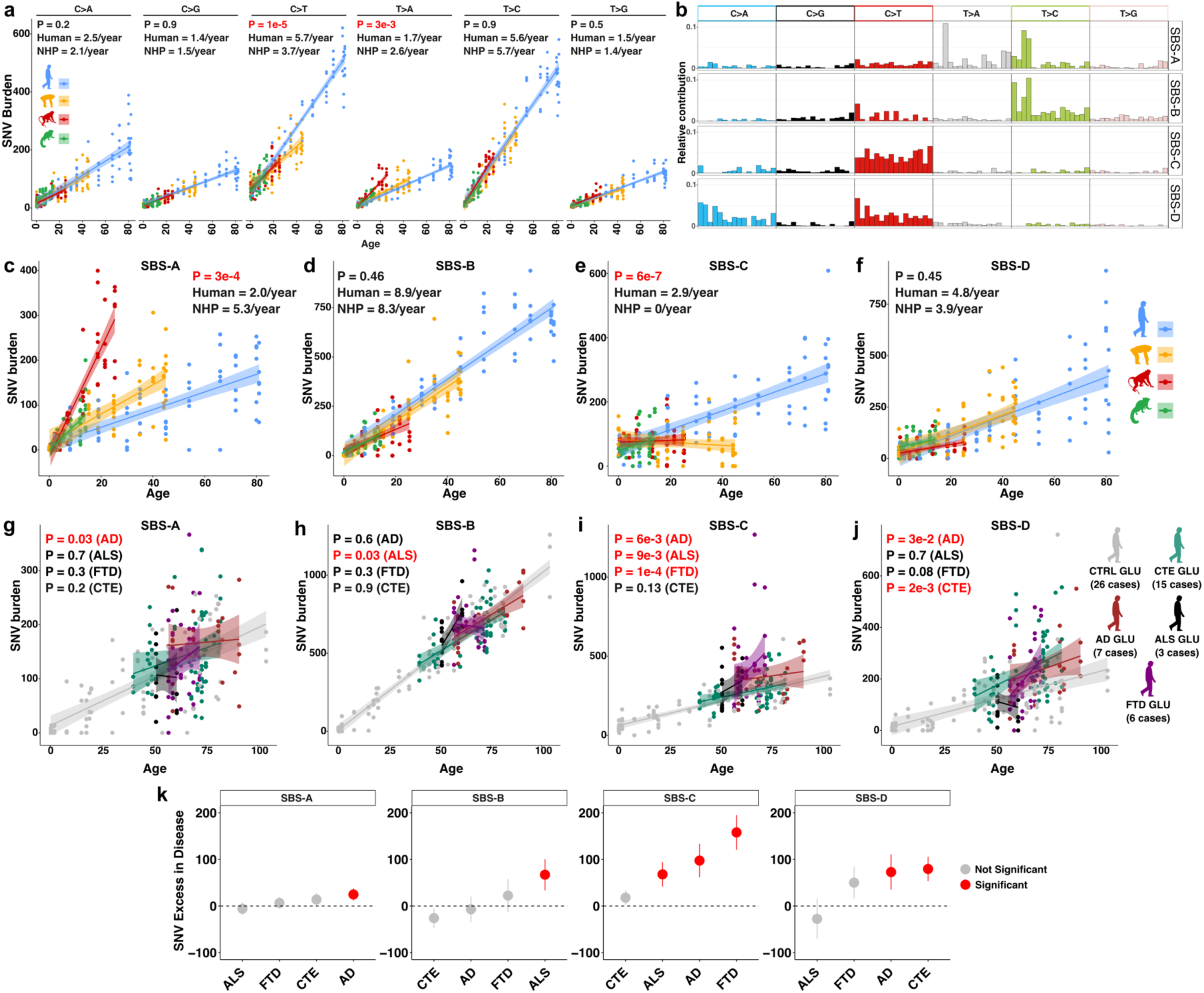
Mutational signatures show divergence across species and neurodegenerative diseases. (a) Genomewide mutation burdens per spectrum in all species for SNVs. NHP: nonhuman primates. Human and NHP mutation rates are calculated using linear mixed model as described in Figure 1. For NHP, the linear mixed model included species as an additional random effect. (b) Profile of de-novo mutational signatures. (c-f) Genomewide mutation burdens of humans and nonhuman primates per mutation signature. Statistics were calculated the as described in Figure 1. (g-j) Genomewide mutation burdens of neurodegenerative diseases per mutation signature. P-values are one-sided and show significance for excess number of mutations in disease compared to healthy control while accounting for age (burden ∼ Disease + Age + (1|Subject)). (k) Average number of mutation difference between disease and control for each mutational signature. Bars indicate standard error of the mean. CTRL: healthy control, CTE: chronic traumatic encephalopathy, AD: Alzheimer’s disease, ALS: amyotrophic lateral sclerosis, FTD: frontotemporal dementia. Significant p-values (P<0.05) are shown in red.

Our results suggest an interesting paradox: that similar rates of mutation accumulation among primates are nonetheless built upon somewhat different patterns of nucleotide substitution, suggesting potentially different mutational mechanisms. We further explored these comparisons among human, chimpanzee and rhesus macaque, excluding marmosets because their low mutation burdens make signatures highly variable. The rate of SBS-A was significantly lower in humans compared to both chimpanzees and rhesus macaques, and SBS-C was significantly higher in humans compared to both chimpanzees and rhesus macaques **(Supplementary Figure 9a)**. Trends towards lower SBS-B and SBS-D rates were seen in rhesus macaques, but did not reach statistical significance **(Supplementary Figure 9a)**. Downward trends of all signatures except SBS-A helps reduce the overall macaque mutation rate to levels comparable to human and chimpanzee mutation rates **(Supplementary Figure 9b)**. Similarly, the human elevation in SBS-C compared to chimpanzees is accompanied by a significant decrease of SBS-A mutations **(Supplementary Figure 9a-b)**. Therefore, despite the similar overall annual mutation rates, the underlying mutational signatures can accumulate at very different rates across species, perhaps indicating an evolutionary constraint on the annual mutation rate, rather than the end-of life mutation burden in neurons (Discussion).

Analyses of cancer datasets have catalogued many mutational signatures and identified the mechanisms underlying some of them, creating a database named COSMIC (the catalogue of somatic mutations in cancer^19^). To gain insight into potential mechanisms underlying our *de novo* signatures, we calculated cosine-similarity between each signature and the COSMIC signatures **(Supplementary Figure 9c-j)** that revealed high similarity (> 0.8) between SBS-B and two signatures, SBS16 and SBS5, as well as between the C>A component of SBS-D and SBS8 **(Supplementary Figure 9d,i)**. The human-specific signature SBS-C is highly similar to SBS30, a signature related to base excision repair pathway^21^, the primary repair pathway for oxidative stress^22^, indicating potentially higher oxidative stress in human neurons **(Supplementary Figure 9e)**^23,24^. In contrast to SBS-B, C and D, none of the COSMIC signatures resembled SBS-A, especially its T>A component **(Supplementary Figure 9c,g,h)**, suggesting that SBS-A reflects a mutational pattern that is rare or absent in human tissues studied to date.

To reveal whether mutational signatures are significantly associated with transcribed regions, we ranked genes based on their transcription level in cortex and tested whether the overlap with each mutation signature is more significant than expected by random sampling of the same trinucleotide contexts **(Methods, Supplementary Table 5)**. SBS-B showed significant association with higher transcription levels in all species, whereas other signatures did not show consistent patterns **(Supplementary Figure 10)**. This is consistent with previous results showing similar transcriptional associations with SBS16, the COSMIC signature that is very similar to SBS-B **(Supplementary Figure 9d)**^25^.

### Evolutionary relationships of mutational signatures associated with neurodegenerative disease

Given the importance of DNA damage and repair for neurodegeneration^8^, we next tested whether human susceptibility to neurodegeneration^9–11^ might be reflected in mutational signatures in normal brains of humans and nonhumans. We analyzed published single-neuron whole genome sequence data from PFC glutamatergic neurons from Alzheimer’s disease (AD), chronic traumatic encephalopathy (CTE), frontotemporal dementia (FTD) and amyotrophic lateral sclerosis (ALS)^26–28^, as well as additional neurons from neurotypical controls^13^, using SCAN2-Zoo. Decomposition of somatic mutation calls with the *de novo* signatures identified in this study revealed that the human-specific signature SBS-C and a conserved signature SBS-D were significantly elevated in at least two degenerative diseases, whereas SBS-A and SBS-B were largely conserved with few exceptions (**Figure 3g-k, Supplementary Table 4)**. Since all four neurodegenerative diseases also display very high levels of 2-4bp deletions (ID4 mutation signatures) as a robust genetic hallmark^26–28^, we also identified COSMIC indel signatures and performed signature decomposition in our data and disease datasets. This analysis recapitulated the known very high levels of ID4 mutations across all neurodegenerative diseases but revealed similar increases of ID4 mutation levels in neurotypical human as well as NHP neurons **(Supplementary Figure 11)**. Taken together, we show that most neurodegeneration related signatures are conserved across primates with the exception of SBS-C, a mostly flat C>T signature that accumulates significantly faster in humans than other primates.

Whereas mutational landscapes of neurodegenerative diseases have previously only been studied in glutamatergic neurons^26–28^ or in neurons without distinction^2^, we find that most signatures accumulate at significantly different rates between glutamatergic and MGE-GABAergic neurons **(Supplementary Figure 12)**. Among the signatures that accumulate faster in neurodegenerative diseases, ID4 and SBS-C also accumulate faster in glutamatergic neurons than in inhibitory neurons, whereas SBS-D accumulates faster in MGE-GABAergic neurons **(Supplementary Figure 12c-e)**. This highlights the importance for elucidating disease associated mutational patterns in neuronal cell type specific manner and could provide further insights into the relationships behind mutational mechanisms and neuronal cell type-specific degeneration.

### Transcriptional alterations in aging human, chimpanzee, and rhesus macaque neurons

The striking lack of evolutionary constraint on end-of-life somatic mutation burden between humans and nonhuman primates indicates that the greatly increased mutation burden in human neurons is unlikely to be fatal in and of itself, yet this higher age-related mutational burden but could still be associated with changes in the transcriptional landscape of neurons. To test this, we isolated and sorted neuronal nuclei from adult and aged subjects across human, chimpanzee, and rhesus macaque dorsolateral prefrontal cortex (**Figure 4a-e**). We matched sample ages across species so that the average age difference between adult and aged samples are ∼40% of the lifespan in all species (**Figure 4b-d**). As detailed in the methods, we combined one individual from each species into a single reaction to mitigate potential tissue processing related artifacts such as ambient RNA contamination^29^. Cells were bioinformatically assigned to species after sequencing by comparing the number of transcripts with zero mismatches to each species’ genome **(see Methods)**.

**Figure 4:**
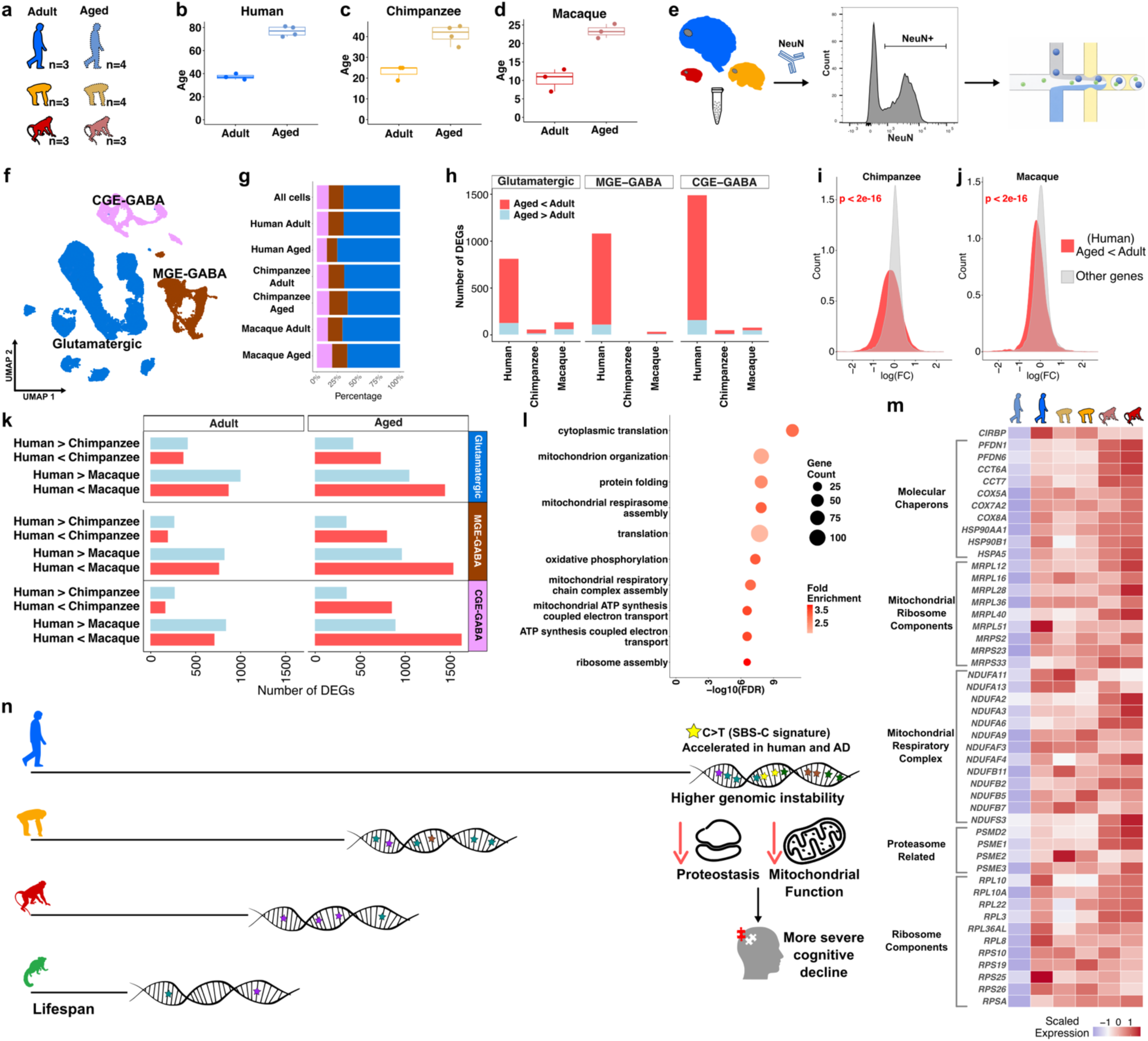
Extensive downregulation of protein synthesis and energy metabolism related genes during aging is human-specific. (a) Number of samples for adult (left) and aged (right) groups per species. (b-d) Age distributions of each sample per species. (e) Schematics showing nuclei isolation, sorting of NeuN+ population followed by single-nucleus RNA-sequencing. (f) Broad annotation of single-nucleus RNA-sequencing result in UMAP space. (g) Breakdown of celltype annotation percentage per species per stage. (h) Number of DEGs (FDR < 0.05 and absolute logFC > 1) between aged and adult individuals in each species. (i-j) Log fold change distributions of genes downregulated in aged individuals in humans (red) and the other genes (gray) in chimpanzee and rhesus macaque DGE results. (k) Number of DEGs (FDR < 0.05 and absolute logFC > 1) between human-chimpanzee and human-rhesus macaque in adult and aged groups. (l) Top ten biological process gene ontologies enriched in downregulated genes in human aging. (m) Heatmap of a subset of genes downregulated in human aging that drive enrichments in mitochondrial and proteostasis gene ontology categories. (n) Summary diagram of the major findings. Increased genomic and cellular instability in human neurons might lead to more severe aging specifically in humans and underlie our unique susceptibility to neurodegenerative diseases.

After quality control, we identified ∼50,000 high-quality neurons and annotated their neuronal subtypes **(Supplementary Figure 13)**, then grouped neuronal subtypes into broader categories matching with the cell type analysis of somatic mutations (**Figure 4f**). Glutamatergic, MGE-GABAergic and CGE-GABAergic (CGE: caudal ganglionic eminence) cell types were detected in all species and stages (adult and aged) with similar proportions (**Figure 4g**). To uncover age-related differential gene expression in all species, we performed differential gene expression (DGE) analysis between adult and aged individuals on the pseudobulk counts in each cell type. Hundreds of genes were significantly downregulated (log_2_FC < -1 and FDR < 0.05) in aged human brain, whereas downregulation was much less common in aged chimpanzee and rhesus macaque, a pattern consistent across cell types (**Figure 4h, Supplementary Table 6)**. Interestingly, genes downregulated in aged humans show similar trends in chimpanzees and rhesus macaques, indicating a similar regulation pattern, albeit with much smaller effect sizes than humans (**Figure 4i-j**). Indeed, lowering the fold change cutoff requirement (absolute log_2_FC > 0.5) revealed many age-related gene expression changes in chimpanzees and rhesus macaques, although downregulated genes were still more pronounced in humans **(Supplementary Figure 14a)**. Quality control metrics such as number of detected genes and intronic read ratio were also similar across species and age groups **(Supplementary Figure 13a-c)**.

We then hypothesized that the extensive gene downregulation in human aging would deviate from the expected pattern of slightly higher levels of upregulated compared to downregulated genes in humans compared to NHPs that has been replicated across studies^30–32^ and performed DGE analysis between species separately for adult and aged groups. We replicated the expected pattern for adults; however, we found hundreds more downregulated genes than upregulated genes in humans compared to both chimpanzees and rhesus macaques (**Figure 4k, Supplementary Table 6)**. A recent study on human dorsolateral prefrontal cortex similarly found extensive downregulation in aged individuals compared to adults^33^ that significantly overlapped with our results **(Supplementary Figure 14b-c)**. In contrast, differentially expressed genes identified in the rodent brain neurons revealed comparable levels of upregulated and downregulated genes **(Supplementary Figure 14d)**^34–36^. These results show that the extensive gene downregulation during aging is uniquely more severe in humans compared to our close relatives and mice.

Gene ontology (GO) enrichment for the genes uniquely downregulated or upregulated in human aging revealed that, while there were no significant enrichments for upregulated changes, nearly all enrichments for downregulated changes were related to proteostasis and mitochondrial functions (**Figure 4l, Supplementary Table 6)**. A total of 804 downregulated genes were in the enriched GO categories, indicating that most age-related gene downregulation in humans reflects decline in proteostasis and energy metabolism. We highlight some of the downregulated genes that function as molecular chaperons, ribosomal proteins, mitochondrial respiratory complex and proteasome related proteins (**Figure 4m**). Additionally, *CIRBP*, a gene highly expressed in long-lived bowhead whales and that likely contributes to its long lifespan by enhancing double-stranded break repair^5^, is >2 fold downregulated uniquely in human aging (**Figure 4m**).

Many studies have shown the age-dependent decline of proteostasis^37^, leading to the conclusion that changes in proteostasis are a hallmark of aging^38^. Age-related declines in energy homeostasis and mitochondrial function are also well documented^34^ and mitochondrial dysfunction is considered to be another aging hallmark^38^. Together, our results indicate that three of the aging hallmarks, genomic instability, loss of proteostasis, and mitochondrial dysfunction are more severe in aged human neurons compared to aged nonhuman primate neurons (**Figure 4m**).

## Discussion

While previous studies have uncovered evolutionary adaptations to sustain genetic integrity in long-living mammalian species^4–7,39^, here we find instead that somatic mutations in neurons accumulate at similar annual rates across mammalian species, resulting in a higher genomic mutational burden in humans compared to other species during aging (**Figure 2**). We further reveal that an underlying mutational signature, mainly composed of C>T transitions, accumulates uniquely in humans and is also elevated across neurodegenerative diseases (**Figure 3**). Finally, we show that extensive gene downregulation during aging is unique to humans and disrupts proteostasis and mitochondrial functions (**Figure 4**). Our results show that longer lifespan may not always be associated with evolutionary adaptations that improve genomic integrity, and that human aging suffers from greater levels of genomic and transcriptomic decline compared to its close relatives.

As the most striking example of a mutational signature that varies between neurons of distinct species, we identified a T>A and T>C dominated signature (SBS-A) that accumulated at very high rates in rhesus macaques, and overall higher rates in NHPs compared to humans. The molecular mechanism of this signature is unknown and could potentially be linked to a DNA repair mechanism that is less error-prone in humans, potentially due to an unknown evolutionary novelty. In contrast, another mutational signature, SBS-C, represented by C>T transitions, significantly accumulates only in humans among the species we studied. While the underlying mutational mechanism for SBS-C is also unknown, it resembles the COSMIC signature SBS30 that accumulates when a base excision repair (BER) enzyme is deficient^21^. Since BER targets oxidative DNA damage, human-specific accumulation of SBS-C might reflect increased oxidative stress-related DNA damage in humans compared to other species, reflecting either intrinsic aspects of neuronal function or perhaps higher levels of neuroinflammation in humans^23,24^. This is also consistent with increased levels of oxidative stress across neurodegenerative diseases^8^ that display a further increase of SBS-C levels compared to neurotypical control neurons.

A previous study on intestinal crypts found that somatic mutation rate is slower in mammalian species with longer lifespan^4^. Given the high cell division and mutation rate of intestinal crypts compared to neurons (∼800/year versus ∼19/year in mouse, respectively), it is possible that tissues composed largely of dividing cells, especially where clonal selection can amplify effects of damaging mutations, may be larger drivers of evolutionary fitness, while somatic mutations in the noncycling neurons of the brain may have more modest fitness consequences because they are not subject to clonal amplification or selection. Indeed, the remarkably similar levels of annual mutation rate across neurons of various species, despite large variations in mutational signatures and hence presumably mutation mechanisms between species, suggests an opposing explanation for neurons, that similar cross-species annual mutation rates indicate a potential evolutionary constraint on the mutation *rate* rather than the end-of-life mutation *burden*.

Conservation of features across species, especially in genomes, frequently implies conserved function, and so it is reasonable to ask whether the conservation of annual mutation rate in neurons of diverse species might implicate conserved functions. At a minimum, the accumulation of 15-20 SNVs per year marks time--more specifically, the age of the individual--with significant precision, and age is a thread around which cognitive experience is structured. Transient DNA damage of several types has been increasingly implicated in synaptic plasticity^40–42^, but whether permanent, double-stranded DNA mutations have some relationship to those plasticity processes has not been tested. Age-related increases in SNV and indels that are enriched at epigenetic sites^13^ could alternatively contribute to normal, age-related reductions in synaptic plasticity in mammals, which could perhaps be evolutionarily constrained. Ultimately, every tissue or indeed perhaps each cell type could reveal its own unique level of evolutionary constraint on somatic mutation rates and burdens, depending on intrinsic mutation rates that might be under evolutionary constraint itself, clonal selection, and contribution to species survival. For example, mouse hematopoietic stem cells accumulate mutations only 3 times faster compared to humans in contrast to 17-fold difference for intestinal crypts^43^, so that neurons are not the first tissues whose annual mutation does not scale with overall organismal lifespan.

Given that SNVs and indels in neurons show only weak to moderate enrichment in functional genomic elements^13,25^, the extensive gene downregulation in aged human brain may not result merely from the greater level of somatic mutation in aged humans. Indeed, we detect no excess of mutational burden in the genes that are downregulated with age in humans compared to the background **(Supplementary Figure 15)**. Other forms of genetic instability such as unrepaired (presumably single-stranded) DNA damage, or genomic repeat length polymorphism^44^ may also accumulate with age, and genetic changes could collectively trigger stress pathways, ultimately resulting in transcriptional downregulation of key biological processes in aged human neurons. In contrast to neurotypical controls, neuronal mutations and specific forms of single-stranded damage appear to reach very high and damaging levels in neurodegenerative disorders^26–28^, some of which may be unique to humans as a species^10,45^. Therefore, somatic mutations in neurotypical controls offer a ‘preview’ of the genetic instability landscape that can be exacerbated in pathological conditions, likely building on the underlying fragility of aging human genomes. Taken together, our results suggest that human aging is more severe than in other species at the molecular level, potentially due to our long lifespan that evolved with little evolutionary constraint to limit the adverse effects of aging on the genome, and that these human-specific aspects of mutational aging are very poorly recapitulated in nonhuman species intended to model the human condition.

## Methods

### Tissue acquisition and maintenance

We obtained fresh frozen postmortem brain tissues from the NIH Neurobiobank (for human tissues), the National Chimpanzee Brain Resource (for chimpanzee tissues) and the National Institute on Aging (RRID:SCR_007324, for rhesus macaque, marmoset and mouse tissues). All tissues were stored at -80°C until the day of the experiment.

Ferret procedures were performed under protocols approved by the Institutional Animal Care and Use Committee at Boston Children’s Hospital. Ferrets (*Mustela putorius furo*) were obtained from Marshall BioResources and were housed in a vivarium under 12 h light / 12 h dark cycle. Food and water were available ad libitum. Ferrets were deeply anesthetized with ketamine (50 mg/ml) and xylazine (20 mg/ml) by subcutaneous injection and transcardially perfused with ice-cold phosphate-buffered saline (PBS) (Thermo Fisher Scientific, Cat. #10010072). Brains were immediately dissected out, placed in a conical tube to be frozen in liquid nitrogen, and transferred to –80°C freezer for long-term storage.

### Nuclei isolation and Fluorescence activated nuclei sorting (FANS)

Postmortem brain tissues were briefly removed from -80° C and small amount of tissue from dorsolateral prefrontal cortex (or frontal cortex for mouse) was collected with a scalpel and transferred to chilled Dounce homogenizer that contained nuclei isolation media (10mM Tris-HCl pH 8, 250mM Sucrose, 25mM KCl, 5mM MgCl2, 0.1% Triton X-100). Tissue was then homogenized with 20-25 strokes of Dounce homogenizer pestle B and lysates were layered on top of a sucrose cushion buffer (1.8M Sucrose, 3mM MgCl2, 10mM Tris-HCL pH 8). To separate the nuclei from cellular debris, the suspension was centrifuged for 40 minutes at 30,000rcf. Supernatant that contained the cellular debris was removed and nuclei were resuspended from the bottom of the tube in a chilled blocking buffer (1X PBS, 0.8% Bovine Serum Albumin) and stained with an anti-NEUN antibody (Millipore MAB377X), anti-SOX10 antibody (Novus NBP2-59621R), and anti-SOX6 antibody (Novus NBP3-44223AF647) for 30 minutes. All antibodies were used at 1:500 concentration. After the antibody staining, nuclei were centrifuged for 5 minutes at 500rcf, and pellets were resuspended in cold blocking buffer. Nuclei suspension was then filtered through a 40uM filter and stained with DAPI (1:1000 of 1mg/ml stock solution) right before FANS.

For FANS, nuclei were gated using size selection and DAPI+ selection followed by antibody gating to isolate specific cell types. For both neurons, NeuN+ signal was gated followed by separation of two populations with SOX6 signal. For glutamatergic neurons, we gated NeuN+ nuclei with lower SOX6 signal, followed by selection of the largest nuclei (top 5%) using the forward scatter. For the MGE-GABAergic neurons, we gated NeuN+ nuclei with higher SOX6 signal. We selected top 50% of this population in terms of both NeuN and SOX6 signals to minimize potential glia contamination that was reported to be up to 5% in NeuN+ population from brain cortex^46^. Using these gates, we then sorted a single nucleus to each well of a 96-well plate for both cell types to be used for single-cell whole genome amplification.

### Single-nucleus RNA sequencing (snRNA-seq) to test sorting purity

To test sorting purity, 17000 nuclei from target population were collected in a 1.5ml tube using an 85µM nozzle. The collection tube contained all components of the reverse transcription (RT) master mix except for RT Enzyme E which was added right before loading the reaction onto the chip. We then followed the GEM generation and barcoding step of the protocol according to manufacturer’s recommendations (10X Genomics, GEM-X Universal 3’ Gene Expression v4). After RT reaction, cDNA amplification and library preparation were performed according to the protocol and libraries were sequenced on an Illumina NovaSeq XPlus sequencer.

### Single-cell whole genome amplification

To perform single-cell whole genome amplification (scWGA) via primary template amplification (PTA), a single nucleus was sorted into a well of a 96-well plate. This was performed for each target population and sample with six replicates on average. PTA was then performed using Bioskryb Genomics Resolve DNA Whole Genome Amplification kit according to manufacturer’s recommendations except that alkaline lysis was performed on ice to prevent potential heat induced artifacts. A negative control reaction that did not contain a nucleus was included in every experiment to test for potential contamination. After PTA, amplified DNA was purified using Ampure beads and the total DNA yield was measured with Qubit. Amplicon sizes of each cell were also assessed with tape-station. Cells with moderate to high (500ng-2µg) and similar yield in each experiment with an amplicon peak > 900bp were then selected for library preparation. 500ng of the PTA product was then used to build the libraries. Libraries were prepared with KAPA Hypreprep according to manufacturer’s protocol except that two double-sided size selection with SPRI (solid-phase reversible immobilization) beads were performed to better enrich for 200-700bp amplicons. Libraries were then sequenced on an Illumina NovaSeq XPlus sequencer.

### scWGA data preprocessing

Raw fastq files were aligned to each species’ own genome using *bwa-mem*^47^. Genome builds used in this study were: *hg19* for human, *panTro6* for chimpanzee, *rheMac10* for rhesus macaque, *calJac4* for marmoset, *mm39* for mouse, *musFur1* for ferret. After alignment, amplification duplicates were marked with *MarkDuplicates* from Picard tools^48^.

To obtain known sites for base recalibration, the default behavior is to use known common variants which is not available in all species. We therefore used the matched bulk WGS for each donor to call single-nucleotide polymorphisms with respect the reference genome and performed base recalibration on all cells by feeding these sites into the *–known-sites* parameter in *GATK BaseRecalibrator*^49^. Base recalibrated BAM files were then directly used to run SCAN2-Zoo.

### Establishing SCAN2-Zoo

To develop SCAN2-Zoo, we modified the source code of SCAN2 with joint genotyping^28^. SCAN2 requires phasing datasets such as EAGLE and SHAPEIT as well as common genetic variants across human populations, both of which are not present and may not be constructed in other mammalians. SCAN2 requires allelic phasing to build an allelic balance (AB) curve from heterozygous single nucleotide polymorphisms (hetSNPs) to statistically model the behavior of putative double-stranded somatic mutations, which displays similar read distributions to hetSNPs^14^. Using AB curve that fluctuates between 0 and 1 throughout the genome, SCAN2’s AB based filters reject a somatic candidate if it can be explained as a pre-amplification artifact (present in one strand; putatively half of the AB) or an amplification artifact (present in less than half of the AB). Since somatic mutations themselves are not phased (i.e we do not know whether they come from one phased allele or the other), it is often possible for a somatic mutation candidate with low variant allele frequency (VAF) to be an artifact arising from one of the alleles. Due to this uncertainty, SCAN2 largely rejects sites with VAF below 0.5^14^.

We reasoned that building an AB curve not from phased alleles but from whichever allele has the highest VAF for hetSNPs could behave similarly to the phased AB curve for somatic mutation candidates greater than 0.5. We implemented this by assigning ‘*phased.hap1*’ the higher count and ‘*phased.hap2*’ the lower count among the alternative and reference counts for each hetSNP that will be used to build the AB. As expected, the resulting AB curve fluctuated between 0.5 and 1 instead of 0 and 1 **(Supplementary Figure 1a-b)**.

We implemented further modifications to SCAN2 to increase its specificity to improve its accuracy across different genomes as detailed below.

1- Requirement for a common genetic variant (e.g dbSNP) filter was removed to accommodate all nonhuman species. We noticed that this filter removes a few calls that are otherwise not filtered as the matched bulk tissue is genotyped as reference homozygous (0/0) by *GATK HaplotypeCaller*. This happens since *GATK HaplotypeCaller* is built for specificity rather than sensitivity. We therefore re-genotyped all candidate calls with *samtools mpileup* in the matched bulk tissue and further removed all candidates with at least one alternate allele supporting read in the bulk tissue.
2- Indel candidates that map to homopolymers that are at least 6nt long were removed as these have a higher risk of being artifactual calls.
3- The site of an indel should contain more deletions than the matched bulk. Any site with many insertions and deletions in the matched bulk could be an artifact of genome alignment. To remove these, we computed the allelic fraction normalized indel fraction difference between the single-cell and bulk as follows.

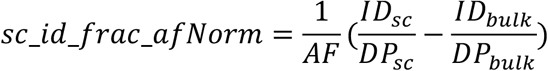

Where AF is the allelic fraction of the indel candidate and ID / DP is INDEL fraction at the site (sc: single cell). Any indel candidates with *sc_id_frac_afNorm* values below 0.5 were removed.
4- For indels, any site with more than two indels (indicating at least two fragments in paired end sequencing) in the matched bulk were also removed.
5- To remove artifactual calls arising from error prone regions during genome alignment (especially in genomes with lower quality), we removed any indel or SNV candidates within 5kb of sites that survive all static filters (i.e filters that do not utilize allelic balance curve) in the same cell.
6- SCAN2 removes indel calls if they are found in at least one other sample in the panel of all other samples but does not apply a panel-based filter for SNVs^13^. We retained the original behavior but also removed any SNVs that are supported by at least 3 reads (i.e at least two fragments in paired end sequencing) in more than half of the samples from other donors in the cross-sample panel.

Since mouse and ferret datasets are smaller in size, we used all samples in the cross-sample panel instead of only non-donor samples. For disease datasets, we used a cross-sample panel of neurotypical human cells to enable direct comparison between disease and neurotypical control cells that were also run with the same panel. Only autosomal chromosomes were used in all datasets, and for ferrets we only used autosomal scaffolds with lengths greater than 50kb. Otherwise, SCAN2-Zoo was applied with the same parameters to all species and disease datasets.

### Genomewide mutation rate, mutation burden estimates

To compare the accuracy of SCAN2-Zoo the original SCAN2 results, we downloaded SCAN2 mutation calls from the published study^13^ and performed SCAN2-Zoo on the same dataset of 52 cells. For accurate genomewide mutation burden estimates in nonhuman species, we used each species own genome size and removed mitochondrial and sex chromosomes as implemented for human genome in the original SCAN2. As quality control, we binned the genome into 50kb intervals and computed MAPD score for each single-cell. MAPD scores higher than 0.5 were not included in mutation rate estimates (however they were included in the cross-sample panel). Aside from MAPD based exclusion, we have also excluded one human donor from neurotypical controls, UMB5823, from all analyses except for the cross-sample panel as further communication with the brain bank revealed that this donor shows neuropathologic changes potentially related to Alzheimer’s disease.

To calculate the genomewide mutation burden for each mutational spectrum (e.g C>T in **Figure 1g**), we computed the scaling factor for each donor by dividing the genomewide mutation burden estimate to total number of mutation calls. Then we counted the total number of mutation calls for each spectrum and multiplied with the scaling factor to computed its genomewide mutation burden estimate.

For all datasets and mutation types (SNVs and indels as well as for individual spectrums and signatures) genomewide mutation rate per year was computed using a linear mixed where subject was fit as a random effect, age in years as a fixed effect and genomewide burden estimate as a response variable using the following formula:

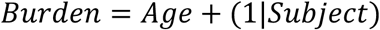

Mutation rate differences between two cell types or species were estimated from their interaction with age as follows:

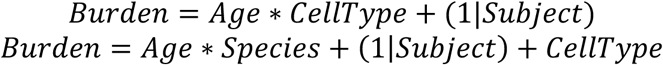

As shown in the bottom formula, cell type information was added to the model as an additional covariate in cases where both glutamatergic and MGE-GABAergic cell types were used.

To calculate the mutation burden estimates by the end of lifespan for each species, we added *genomewide mutation rate X* (*species lifespan* − *donor age*) to each cell’s genomewide mutation burden estimate. We used the species lifespan as reported in studies that observed each species in captivity^50–54^ (chimpanzee: 39 years, rhesus macaque: 27 years, marmoset: 12 years, mouse: 2.75 years, ferret: 7 years). For humans, we used 82 years as estimated in a previous study^4^.

### Multiplexed end-tagging amplification of complementary strands (META-CS)

Tn5 transposition, strand tagging, PCR, and library prep was performed as previously described, but with minor modifications^17^. In short, Tn5 transposomes were added to nuclei for transposition as before. In our case, enzymatic deactivation of the Tn5 transposase was done with 0.05 uL NEB Themolabile Proteinase K per sample incubated for 37°C for 30 min, then 55°C for 10 min, followed by a 4°C hold. Strand tagging was performed as before, but degradation of strand-tagging primers was performed with 1 uL of NEB thermolabile Exonuclease, incubated for 37°C for 15 min, 65°C for 5 min, and then held at 4°C. Samples were indexed and PCR-amplified using premixed NEB index primers and only 11 PCR cycles to minimize amplification artifacts. PCR products were purified using Zymo DNA Clean & Concentrator columns by adding 200 µL binding buffer to 50 µL reaction volume, following the manufacturer’s protocol with an additional centrifugation step after removal of the wash buffer. DNA was eluted in 50 µL Low TE buffer. Ampure bead size selection was performed to isolate ∼450 bp fragments for sequencing.

### Meta-CS Data Processing Pipeline

META-CS data was processed and somatic SNVs were called using a customized pipeline that integrates elements from pre-pe (https://github.com/lh3/pre-pe) and lianti (https://github.com/lh3/lianti), with substantial modifications to support multi-allelic input. A key innovation was the introduction of barcode extraction and pooled read handling. Reads entering pileup were required to meet quality thresholds (minimum mapping and base quality of 30). A variant was called if supported by ≥8 total reads (-a8) with at least four reads from each strand (-s4). Filtering criteria excluded: (i) sites within 100 bp of each other, and (ii) variants within 10 bp of read ends. The procedure identified all ALT alleles, including germline SNPs and somatic SNVs; ALT alleles without bulk read support were classified as somatic. False negative rates for SNV detection were estimated by comparing germline heterozygous SNPs called in duplex data with those observed in the bulk. The number of haploid genomes was estimated by calculating the weighted average of unique molecules across germline heterozygous SNP (gHetSNP) sites, using the following formula:

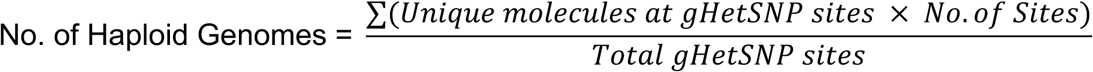

The mutation rate was then adjusted per genome by dividing the total number of detected duplex sequencing SNVs by the estimated number of haploid genomes:

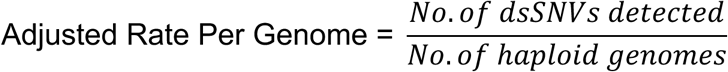

### De-novo mutational signature extraction

De-novo mutational signature extraction benefits from large number of samples to obtain clearly segregated signatures. Our entire dataset could be used for this analysis, however different genomes may have different background (i.e opportunity space) for mutational contexts that can skew the results. Therefore, we first calculated the ratio of each trinucleotide context in the genome for all species and correlated the relative ratio of all trinucleotide contexts to assess similarity of the opportunity space. The correlation was nearly perfect among primates but lower between human and mouse **(Supplementary Figure 16, Supplementary Table 7)**. Based on this result, primate cells from all neurotypical donors were used in extracting de novo mutational signatures.

To obtain robust results, non-negative matrix factorization (*nmf*) from the R package *MutationalPatterns*^55^ was run with 200 random initializations. We extracted 4 de-novo signatures as the cophenetic correlation coefficient across the random initializations was stable up to 4 signatures. To find the cosine similarity with known signatures, we downloaded COSMIC SBS signature database (v3.3.1) for the GRCh37 genome build and calculated the cosine similarity between each de-novo signature and all COSMIC signatures.

To fit de-novo signatures to disease datasets, we performed nonnegative least squares regression using *fit_to_signatures* from *MutationalPatterns* package and calculated the genomewide mutation burden estimate of each signature as described above.

### Signature decomposition of indels

Given the sparsity of indel calls in neurotypical controls, we relied on identifying the contribution of known indel signatures rather than de-novo mutation extraction. To do this, we used *SigProfileExtractor* on human results to identify active mutational signatures (COSMIC v3.3.1). This yielded ID4, ID8, ID9 and ID11 as active signatures. We then performed nonnegative least squares regression and calculated the genomewide mutational burden of each signature as described before. This was performed for all species as well as neurodegenerative diseases.

### Enrichment of mutational signatures in transcribed genes

Testing statistical enrichment of SNV calls that overlap with a custom set of genomic intervals, such as transcribed regions, requires randomized backgrounds that are selected with similar constraints that apply to SNV calls. Since SNV calling is only possible in regions with certain depth, we first generated a set of callable regions that have at least 10 reads in the *SCAN2-Zoo* output after genotyping step (*joint_depth_matrix.tab.gz* files). We then randomly subset the callable regions to 200,000 sites, and for each mutation signature we randomly selected sites from the callable regions based on each signature’s trinucleotide contexts such that the number of sites per trinucleotide context is proportional to the probability of that trinucleotide context in the given mutation signature’s spectrum. The number of random sites per trinucleotide context per mutation signature was matched to the number of called SNVs for the same trinucleotide context in the same mutation signature. Randomized sites were selected 1000 times to create a null distribution for each mutational signature.

To obtain a list of genes divided by their expression levels in the prefrontal cortex, we downloaded GTEX dataset^56^ (*gene_tpm_v11_brain_frontal_cortex_ba9.gct.gz*) and divided the genes into 3 equal groups based on their TPM (transcript per million) normalized expression values. The gene coordinates were extracted from each species’ own gene annotation file that were linked to the genome builds used in this study (see *Data Preprocessing*). Number of overlapping sites were counted for SNVs and the 1000 matched randomized background for each mutational signature. We then calculated the empirical p-value as the ratio of randomized sites that break the expectation (i.e p=0.01 for enrichment indicates that 10/1000 randomized sites had a better overlap with the given genomic regions than the observed SNV calls). P-values < 0.01 were considered as significant enrichment or depletion.

### snRNA-seq for aging comparisons

We followed the same steps described previously in “Single-nucleus RNA sequencing to test sorting purity**”** except that we sorted 5660 NeuN+ nuclei from one human, one chimpanzee and one rhesus macaque tissues into the same 1.5ml conical tube. All tissues were from the dorsolateral prefrontal cortex and post-mortem. Collecting different species (or even samples with different genotypes) into the same reaction with subsequent bioinformatic cell to species assignment should also reduce potential ambient RNA contamination imbalance that might be observed in tissues from different resources^30^ since all tissues are processed in the same reaction. We sequenced all libraries at a high depth (125G/sample) on an Illumina NovaSeq XPlus sequencer (25B flow cell, 2×150bp).

### snRNA-seq analyses to test sorting purity

Fastq files were preprocessed with *cellranger count*^57^ using each the reference genome for each species (human: hg19, chimpanzee: *panTro6*, rhesus macaque: *rheMac10*, marmoset: *calJac4*). We then calculated intronic read ratio using *dropletQC*^58^ for all cells that are called by *cellranger count* and used the *R* package *Seurat*^59^ for the subsequent analysis. Following normalization (*SCTransform*), dimensionality reduction by principal component analysis (PCA) and clustering at high resolution (resolution=1), we removed any clusters with low intronic read ratio compared to other clusters (typical median value was <40%) as these are unlikely to represent intact nuclei. We then plotted known cell type markers of all major brain cell types for all clusters and calculated the sorting purity as the percentage of cells that only expressed the given cell type’s markers.

### Preprocessing of snRNA-seq data on aging

To account for potential gene annotation quality differences between species, we performed liftoff^60^ of *hg19* gtf to both *panTro6* and *rheMac10* assemblies and built cellranger references with these gtf’s using *cellranger mkfastq*. Since all libraries contain cells from more than one species, raw reads in fastq files were mapped to all three species (human: *hg19*, chimpanzee: *panTro6*, rhesus macaque: *rheMac10*). Raw count matrices from the cellranger count output were used to remove ambient RNA contamination with *CellBender*^61^ in each species. Cells called by CellBender were then assigned to their species as described in the next section. Dimensionality reduction, clustering and quality control was performed separately for each species as described in the previous section. Potential doublet clusters that have high levels of known cell type specific marker gene expression were removed. Small percentage of glia contamination that is expected in NeuN+ sorting^46^ was also removed. Cells across species were then integrated using Seurat’s reciprocal PCA integration method^59^. To annotate cell types, SEA-AD dataset from Allen Institute for Brain Science was used as the reference dataset and label transfer was performed using Seurat. Each cluster was assigned a label based on the dominant cell type annotation to that cluster. Resulting fine cell type annotation was then grouped into broader categories as glutamatergic, MGE-GABAergic and CGE-GABAergic neurons based on marker gene expression and prior knowledge^62^.

### Cell to species assignment in snRNA-seq data on aging

For cell to species assignment, we reasoned that mismatches should be frequent between all species given that even human and chimpanzee sequences differ, on average, 2 in every 100 base pair. Therefore, if only the perfectly matching reads are counted, a cell that comes from human tissue would have substantially more UMIs (unique molecular identifiers) if aligned to a human genome as opposed to a chimpanzee genome, enabling a straightforward assignment to species. We implemented this strategy in following steps:

1- For each species alignment, BAM file was subset to retain reads with no mismatch.
2- BAM file was further subset to retain only the cells called by *cellranger count*.
3- Gene to cell matrix was generated with *umitools count*^63^ from this BAM file.
4- For each cell, species assignment was performed by using the following linear mixed formula model in *R*: *umi count* = *species* + (1|*genes*) to test significance of species contribution while accounting for gene-to-gene variability. This was performed first for rhesus macaque and others, then for human and chimpanzee for the remaining cells.

For cells assigned to humans, perfectly aligned UMI count in humans was ∼3 fold more than chimpanzees and ∼32 fold more than rhesus macaques **(Supplementary Figure 17)**. We observed similar patterns for cells assigned to chimpanzees and rhesus macaques with perfectly aligned UMI counts being at least several folds greater in the assigned species compared to other species **(Supplementary Figure 17)**. DGE analyses (explained below) also yielded several fold greater number of DEGs between human and rhesus macaque compared to between human and chimpanzee consistent with previous studies^30–32^.

### Differential Gene Expression

Orthologous genes across human, chimpanzee and rhesus macaque were identified using the R package *orthogene*^64^. All gene expression matrices were subset to contain only the orthologous genes. For each cell type and comparison (either adult vs aged or between species) lowly expressed genes (present in <1% of all cells) were removed. UMI counts were summed per gene across all cells of each individual to generate the pseudobulk matrix which was then used as input to *edgeR*^65^ to statistically test the differential expression for each gene. Genes with FDR < 0.05 and absolute log_2_FC > 1 were determined to be statistically significant.

### Enrichment analyses

DEGs that are downregulated or upregulated in human aging, but not in chimpanzee or rhesus macaque aging, in at least one cell type were tested for gene ontology enrichment using *clusterProfiler*^66^. All genes tested for differential gene expression were used as the background gene list. Significant enrichment was determined as those with p.adjust < 0.01.

To compare aging downregulated genes in this study and Jeffries et al^33^, genes downregulated in at least one cell type were pooled per study. Then the statistical significance of the overlap between studies was evaluated with p-value obtained from Fisher’s exact test.

### Mutation burden comparisons for gene sets

Genomic regions for all human SNVs and indels were extracted and combined as a single list. Then genomic coordinates for all genes were extracted and expanded 50kb upstream to increase the level of overlap with mutations and account for the mutations overlapping potential regulatory regions. Raw mutation counts were then normalized to gene length (plus 50kb) for each gene set (non significant background genes, all aged < adult DEGs and aged < adult DEGs with mitochondria or proteostasis related functions as determined by their presence in the gene ontology terms with these functions.

### Interactive visualization of snRNA-seq data

The processed snRNA-seq object was reduced in size using DietSeurat^67^ by retaining only the normalized expression data and UMAP dimensional reduction, while removing raw count matrices and intermediate data structures. This step was performed to satisfy ShinyApps.io deployment constraints on application size. An interactive web application was generated using the ShinyCell framework. Configuration files were created using createConfig, and application components were generated using makeShinyFiles. The resulting Shiny application was deployed to the ShinyApps.io platform, enabling browser-based exploration of cell populations, metadata, and gene expression patterns. The interactive dataset is available at: https://walshlab-bch.shinyapps.io/shinyApp/.

## Acknowledgments

We thank Dr. Alice Li for help with human brain tissues and other members of the Walsh lab for useful discussions. We also thank the BCH FACS Core Facility for equipment use and assistance. E.C. is supported by Jane Coffin Childs Fellowship from the Howard Hughes Medical Institute. This research was made possible in part using biomaterials from the NIA Nonhuman Primate Tissue Bank (https://www.nia.nih.gov/research/dab/nonhuman-primate-tissue-bank) at the Wisconsin National Primate Research Center, University of Wisconsin-Madison under contractual agreement with the National Institute on Aging (NIA) as well as from the NIA Aged Rodent Tissue Bank (Aged Rodent Tissue Bank | National Institute on Aging (<u>nih.gov</u>) at the University of Washington, Seattle under contractual agreement with the National Institute on Aging (NIA), and the National Chimpanzee Brain Resource (www.chimpanzeebrain.org; supported by NIH grant NS092988). This work was supported by funding from the Allen Family Philanthropies and Grant 62587 from the John Templeton Foundation (the opinions expressed in this publication are those of the author(s) and do not necessarily reflect the views of the John Templeton Foundation) to C.A.W.. The National Institutes of Health (NIH) supports C.A.W. (NINDS grant R01NS032457, NIA grant R01AG070921, and SMaHT Consortium grant UG3NS132138), to P.J.P. (SMAHT Consortium grant UH3NS132138), and C.C.S. (NIA grants R01AG067419 and R01AG087945, NIMH grant R01MH134809, and NHGRI grant R01HG011641). C.A.W. is an Investigator of the Howard Hughes Medical Institute. Human tissues were obtained from the NIH Neurobiobank at the University of Maryland, Baltimore, MD and at the NIH Brain & Tissue Repository-California, Human Brain & Spinal Fluid Resource Center, VA West Los Angeles Medical Center, Los Angeles, CA, which is supported in part by the National Institutes of Health and the US Department of Veterans Affairs.

## Competing interests

P.J.P. is a member of the scientific advisory board (SAB) for Bioskryb Genomics, Inc. C.A.W. is a member of the SAB of Bioskryb Genomics, Inc, (cash, equity), Mosaica Therapeutics (cash, equity), and an advisor to Maze Therapeutics (equity).

## Data and Code availability

Previously published sequencing data were downloaded from dbGaP. Sequencing data generated in this study will be deposited in a public repository. For human data, sequencing data will be deposited under controlled-access conditions in compliance with human subject privacy regulations. SCAN2-Zoo will be made available through GitHub. All other scripts and code will also be made publicly available on Zenodo and GitHub.

