## Supplementary Figures for "Somatic mutations and single-nucleus transcriptomics reveal uniquely human properties of neuronal aging"

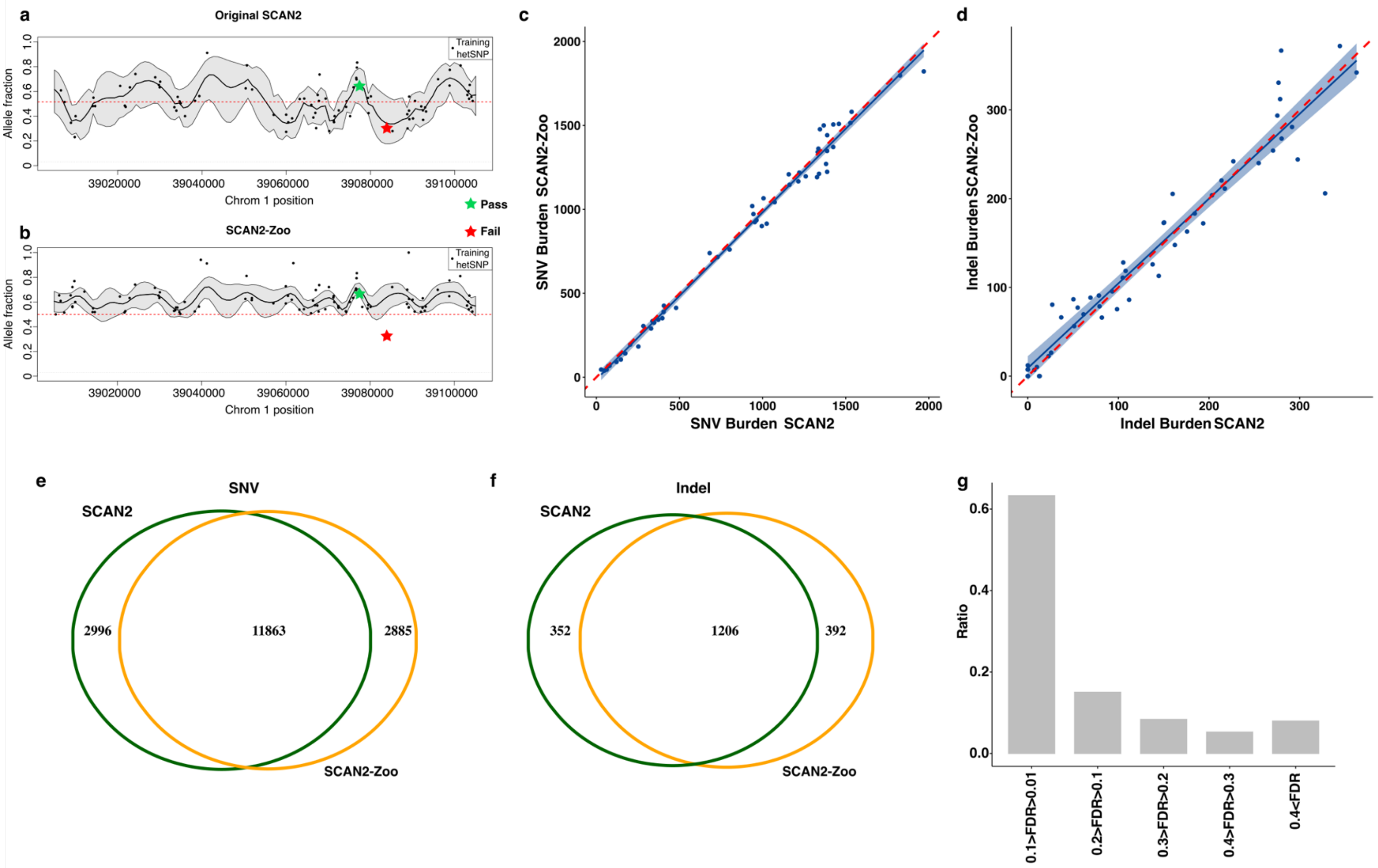


**Supplementary Figure 1: Comparison of the original SCAN2 and SCAN2-Zoo mutation burdens.** (a-b) Allelic balance of the same region in the (a) original SCAN2 and (b) SCAN2-Zoo. Each dot represents a heterozygous SNP that is used to build the allelic balance curve. The red star indicates a somatic mutation candidate that would fail both the original SCAN2 and SCAN2-Zoo criteria. This candidate would have a high chance of failing the original SCAN2 criteria because even though its allele frequency is on the allelic balance curve and it could be a true somatic mutation, it could also be an artifact from the other copy of the chromosome since its allelic frequency is below 0.5. In other words, since the other copy's allelic balance is ~0.6, an artifact from one strand could have a VAF around 0.3 (assuming even strand amplification). Similarly, this somatic mutation candidate would also fail SCAN2-Zoo since it is not on the allelic balance curve built by the heterozygous SNP with maxVAF and would be interpreted as a potential artifact (see Methods). In contrast, somatic mutation candidate depicted by the green star would survive the filter in both tools as its VAF is greater than 0.5 and it is on the allelic balance curve. (c-d) Mutation burdens of glutamatergic neurons across ages in the original SCAN2 (x-axis) and SCAN2-Zoo (y-axis) for (c) SNVs and (d) indels. Shaded blue shows 95% confidence interval and dashed red line shows perfect agreement (x=y). Each data point represents a single-cell. Results for "SCAN2 Original" are taken as-is from the original study that developed SCAN2[14]. (e-f) Overlap of the mutation calls between the two methods for (e) SNVs and (f) indels. (g) Many of the mutations found in SCAN2-Zoo but not in the original SCAN2 fails due to the stringent pre-amplification artifact filter (*lysis.fdr* in SCAN2). This plot breaks down *lysis.fdr* value of such sites into brackets (x-axis) and shows the ratio of each bracket among all such sites (y-axis).

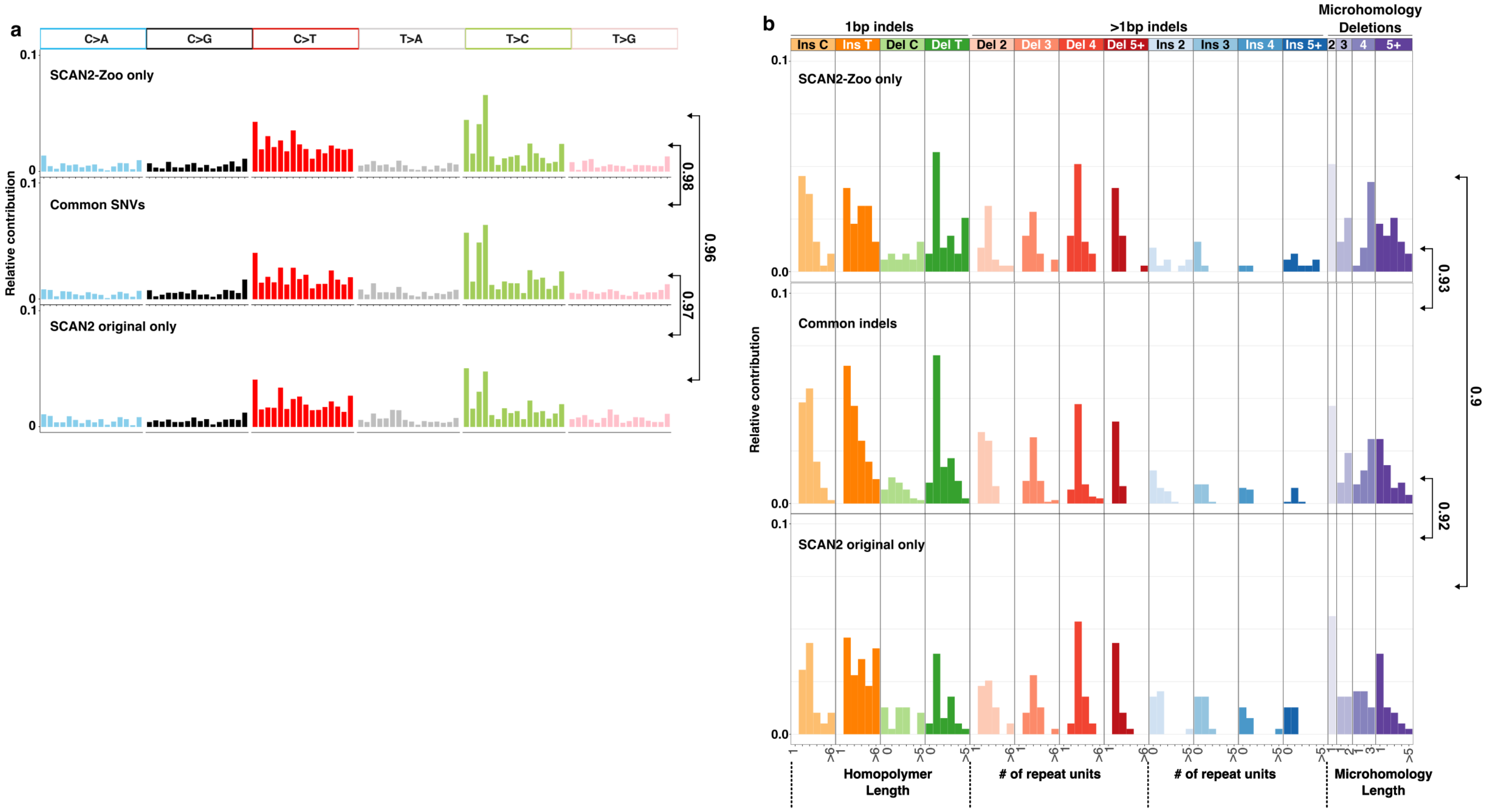


**Supplementary Figure 2: Comparison of the original SCAN2 and SCAN2-Zoo mutation spectrums.** (a) Plots display the mutational spectra of SNVs found only in SCAN2-Zoo (top) or commonly found in both SCAN2-Zoo and the original SCAN2 (middle) or found only in the original SCAN2 (bottom). (b) Same as panel a but for indels. Numbers on the right indicate cosine similarity between the indicated pair.

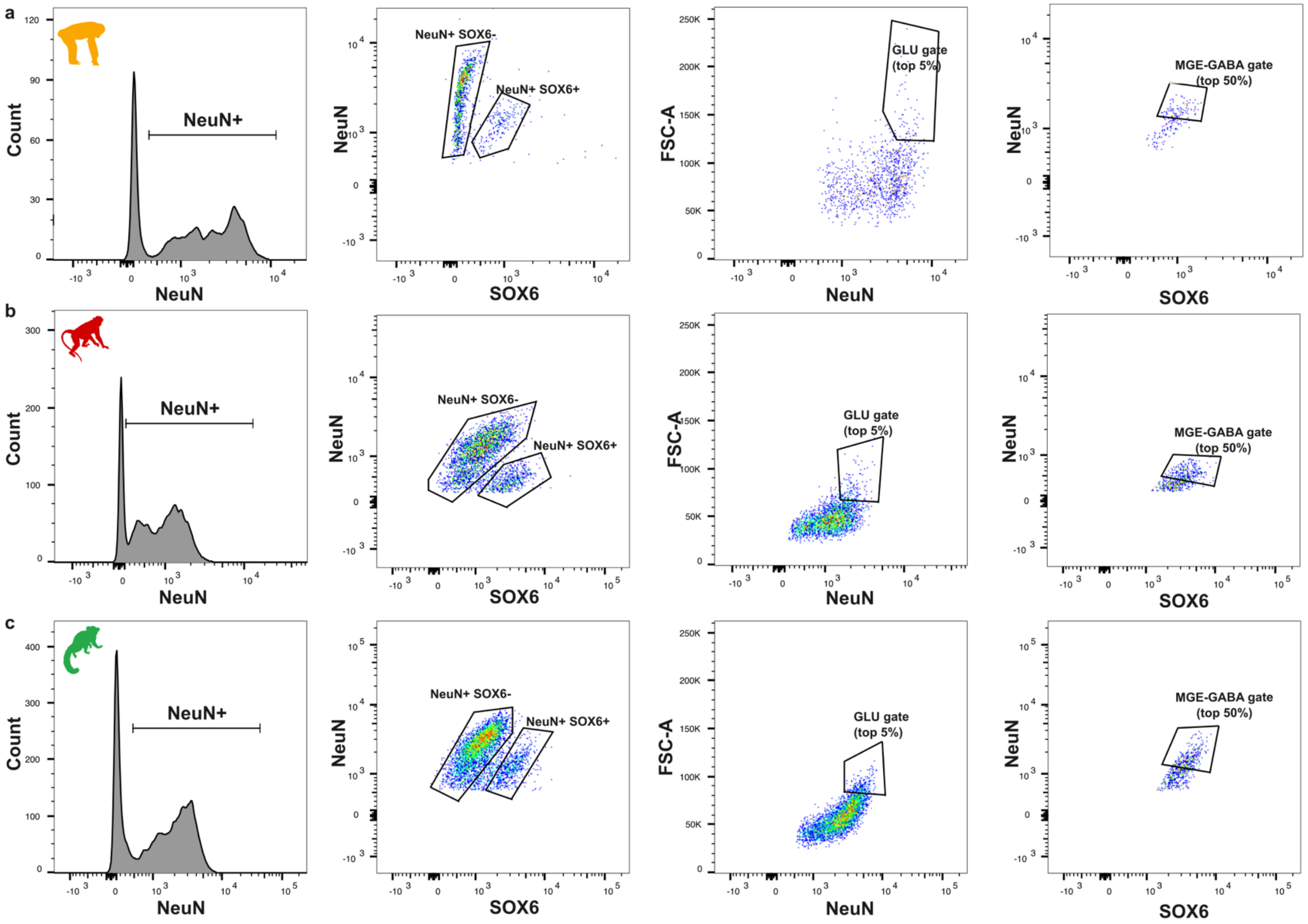


**Supplementary Figure 3: Fluorescence activated nuclei sorting strategy in NHPs.** Sorting gates follow the same strategy as described for humans in Figure 1a. (b) shows Chimpanzee sample 6237, (c) shows rhesus macaque sample 413, (d) shows marmoset sample 446. Figures were generated using FlowJo version 10.

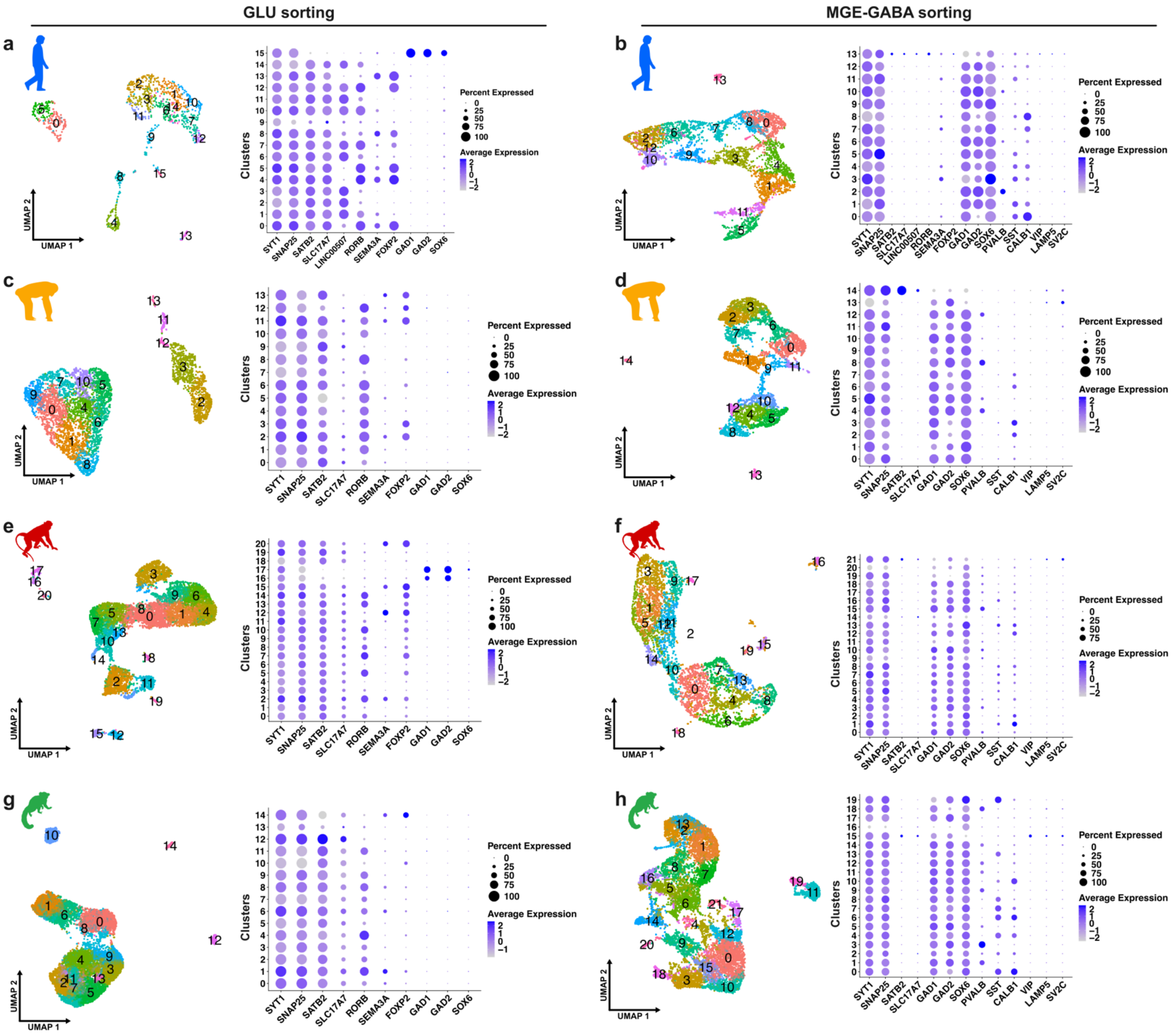


**Supplementary Figure 4: Sorting purities tested by snRNA-seq.** (a-b) snRNA-seq results of nuclei sorted from the (a) 'GLU' or (b) 'MGE-GABA' gates described in Figure 1a and Supplementary Figure1. Clusters on UMAP are described on the left of each panel. Relative transcript levels of marker genes for each cluster are shown on the right of each panel. Numbers on plots indicate clusters. (c-d) Same as in a-b but for chimpanzees. (e-f) Same as in a-b but for rhesus macaques. (g-h) Same as in a-b but for marmosets.

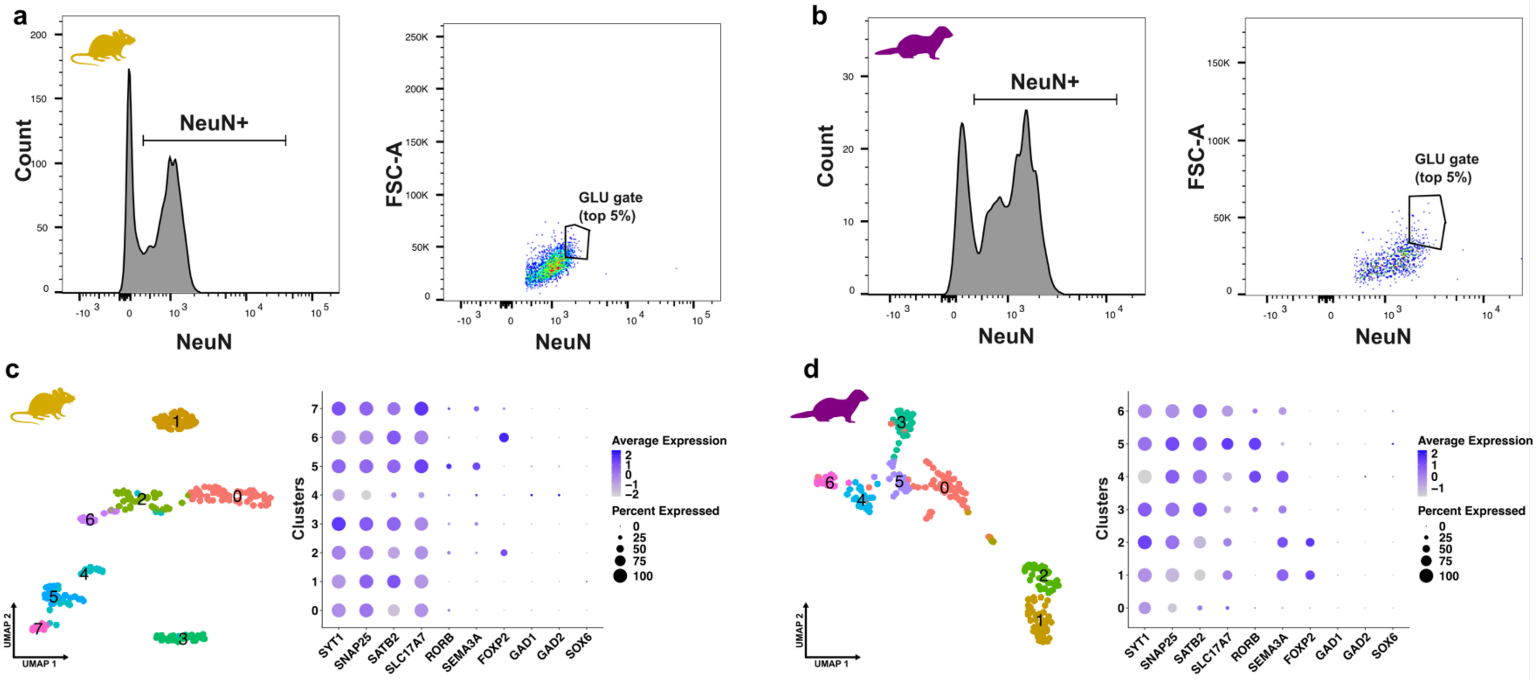


**Supplementary Figure 5: Fluorescence activated nuclei sorting and sorting purities of mouse and ferret glutamatergic neurons.** (a-b) (a) Mouse and (b) ferret Sorting gates follow the same strategy as described for humans in Figure 1a except for the separation on MGE-GABA neurons by SOX6 staining. (c-d) snRNA-seq results of nuclei sorted from the 'GLU' gates for (c) mouse and (b) ferret. Clusters on UMAP are described on the left of each panel. Relative transcript levels of marker genes for each cluster are shown on the right of each panel. Numbers on plots indicate clusters.

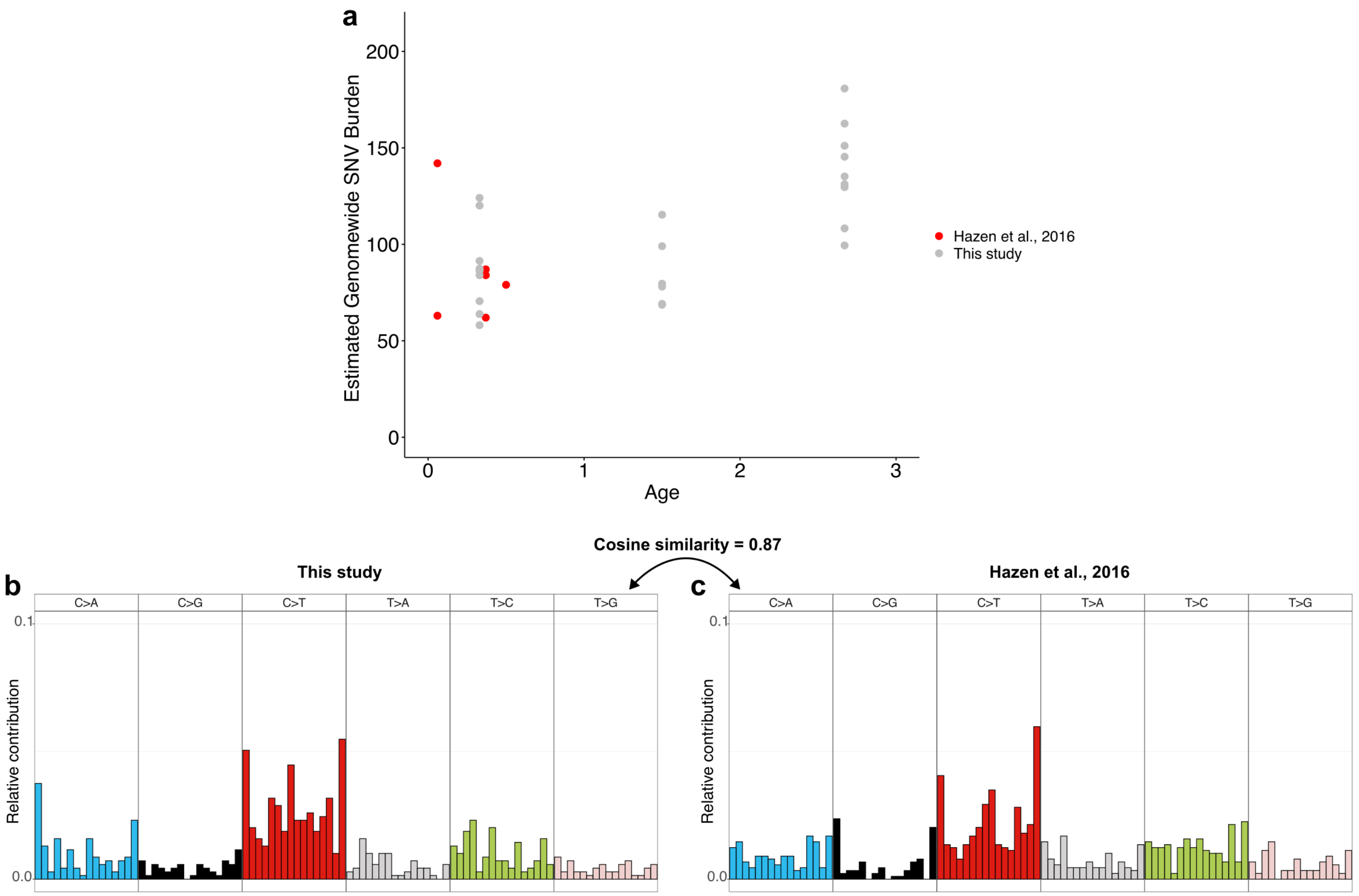


**Supplementary Figure 6: Comparing the similarity between mouse neuron SNVs in this dataset and Hazen et al.** (a) Mutational spectrum of all SNV calls in this dataset. (b) Mutational spectrum of all SNV calls in Hazen et al.[15]. (c) Estimated genomewide mutational burden in both datasets. Each datapoint indicates a cell for this study and a sample derived from a single-cell in Hazen et al. All results from Hazen et al. are displayed without any further analysis and reflect the original results.

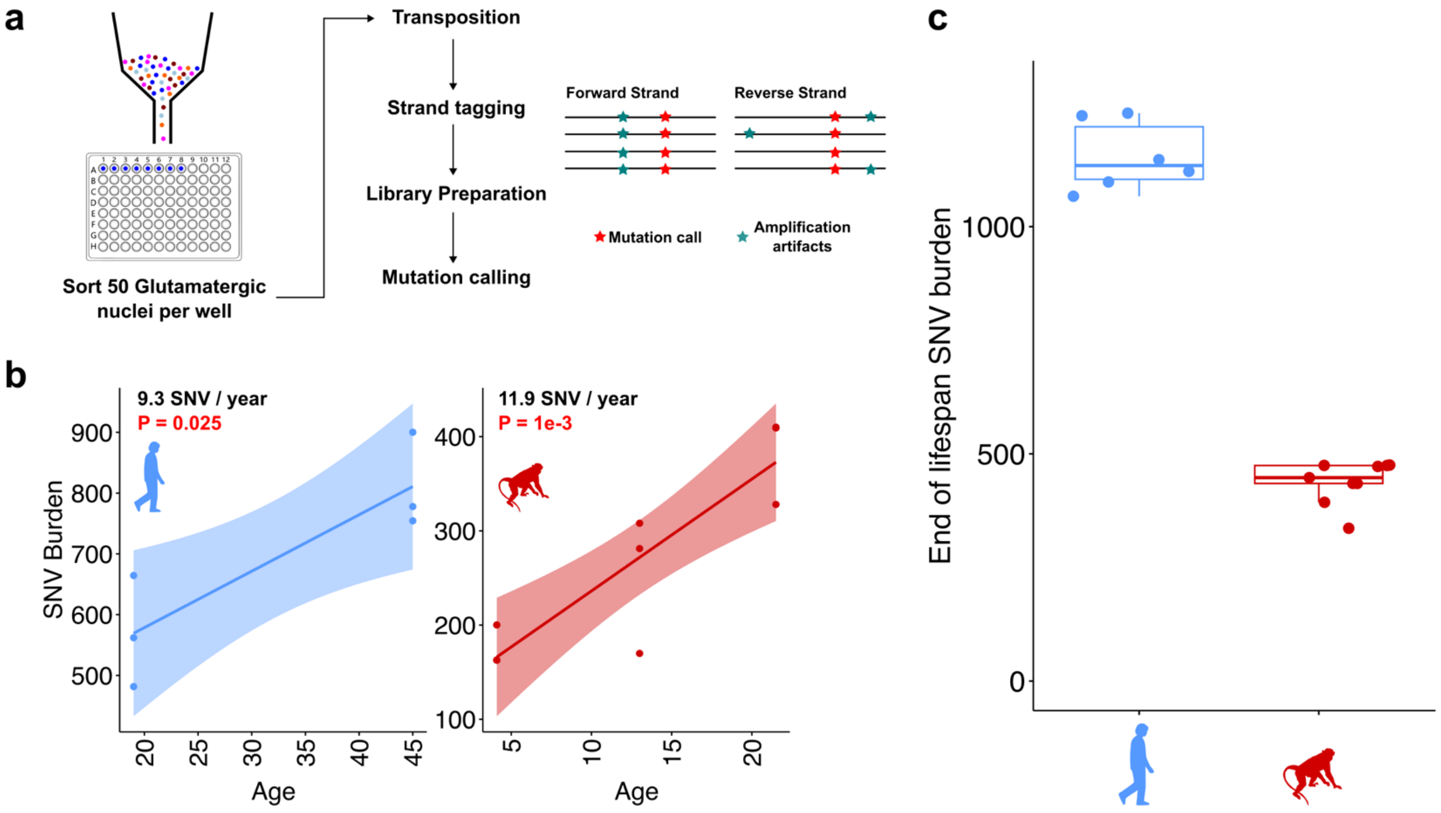


**Supplementary Figure 7: META-CS results on human and rhesus macaque glutamatergic neurons.** (a) Experimental outline of META-CS. Each META-CS sample contains 50 glutamatergic neuron nuclei sorted with the same strategy described in Figure 1a. META-CS involves transposition and strand tagging to distinguish amplification and sequencing artifacts that are not present in both strands from true somatic mutation calls that are present on both strands. (b) Genomewide mutation burden estimates in human (left) glutamatergic neurons and (right) rhesus macaque glutamatergic neurons. (c) End of lifespan burden estimates of human and rhesus macaque samples. All statistics were computed using the same approach described in Figure 1 and Figure 2.

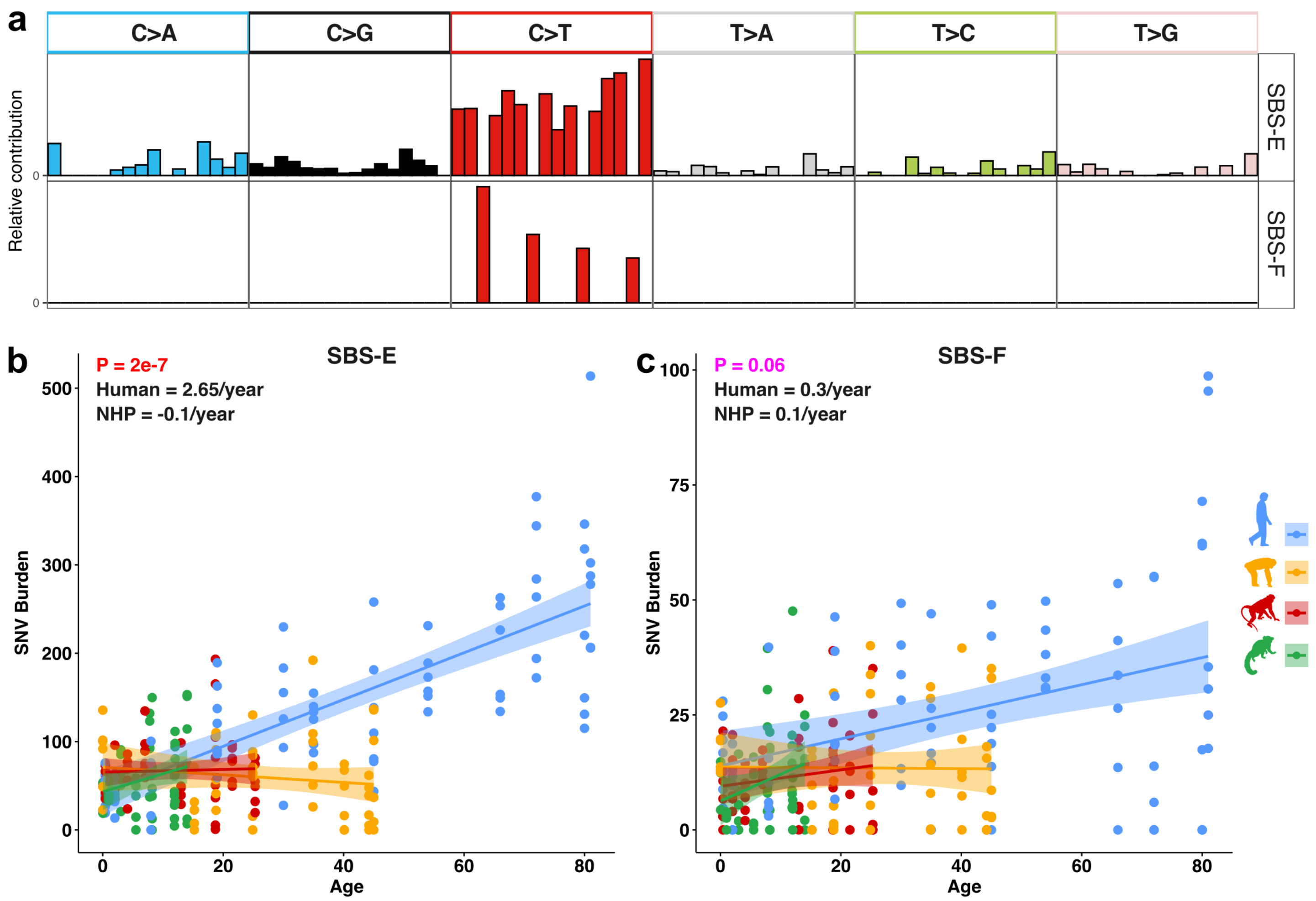


**Supplementary Figure 8: Breakdown of SBS-C into SBS-E and SBS-F based on C>T at CpG sites.** (a) Mutation spectra of SBS-E and SBS-F. SBS-E is C>T at CpG sites removed from SBS-C and SBS-F is the C>T at CpG sites from SBS-C. (b-c) Genomewide mutation burden estimates of (b) SBS-E and (c) SBS-F across species. Statistics are calculated as described in Figure 1.

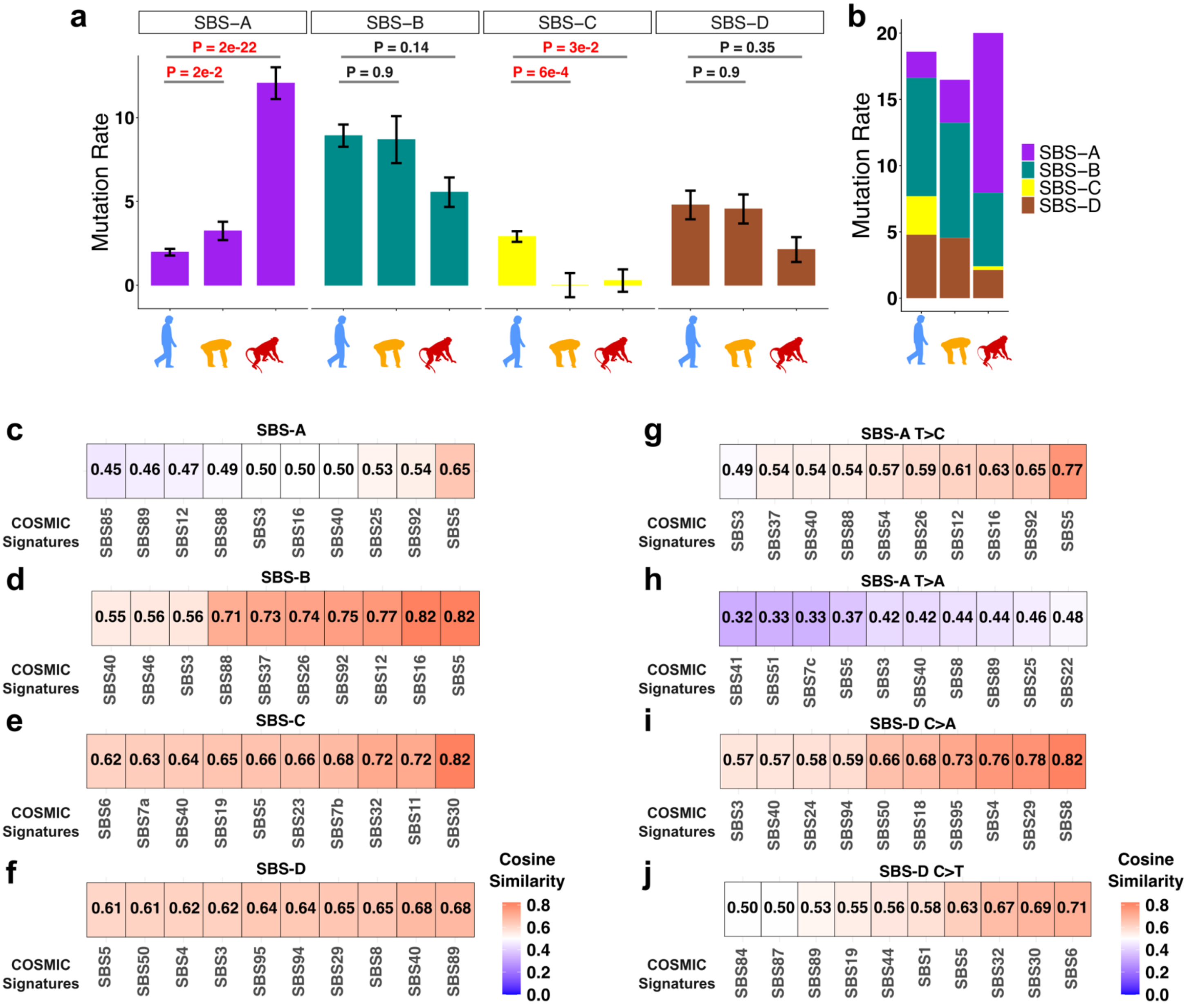


**Supplementary Figure 9: Further comparisons on mutation signatures.** (a) Mutation rate per year for each mutational signature in human, chimpanzee and rhesus macaque. P-values are calculated by testing the contribution of species difference in the to the mutation rate using the model *burden ~ Species*Age + (1|Subject) + CellType*. (b) Breakdown of mutation rates by signature in each species. (c) Cosine similarities between COSMIC mutational signatures and SBS-A. Top 10 COSMIC signatures with the highest similarities are shown. (d-f) Same as panel c but for SBS-B, SBS-C and SBS-D. (g-h) Same as panel c but only for (g) T>C spectrum or (h) T>A spectrum. (i-j) Same as panel f but only for (i) C>A spectrum or (j) C>T spectrum.

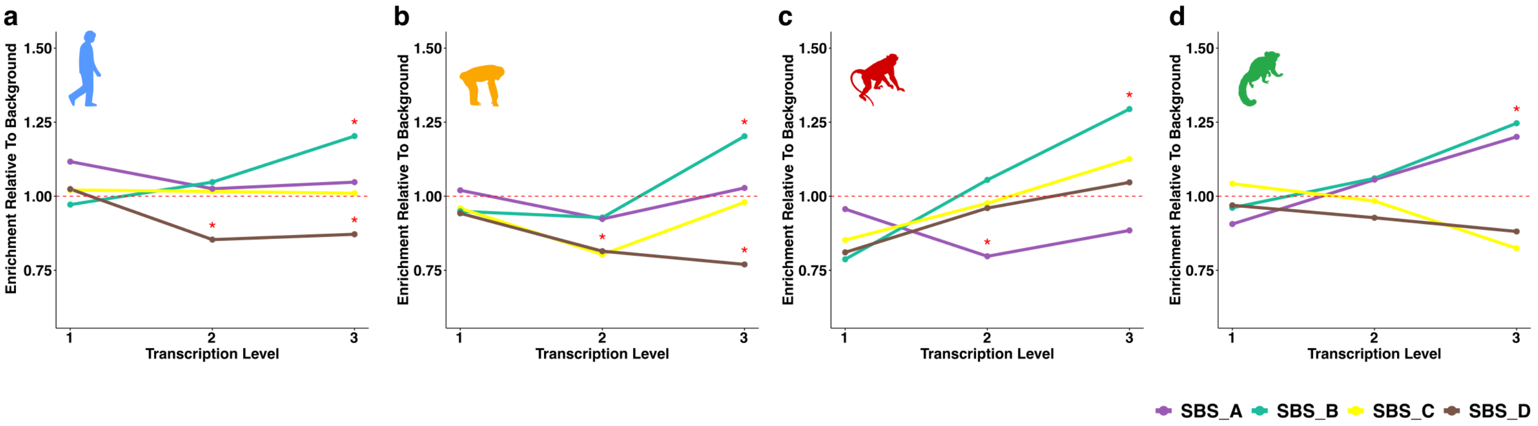


**Supplementary Figure 10: Enrichment of mutational signatures in gene groups separated by their transcriptional level.** Genes were split into three groups based on transcription levels (transcript per million normalized gene expression values of dorsolateral prefrontal cortex in humans (see Methods)). Levels 1, 2, 3 indicate the low, medium, high transcription levels respectively. Asterisks indicate significant enrichment ($p < 0.05$) relative to background as detailed in the Methods. Panels a, b, c, d show enrichment results per mutational signature for human, chimpanzee, rhesus macaque and marmoset, respectively.

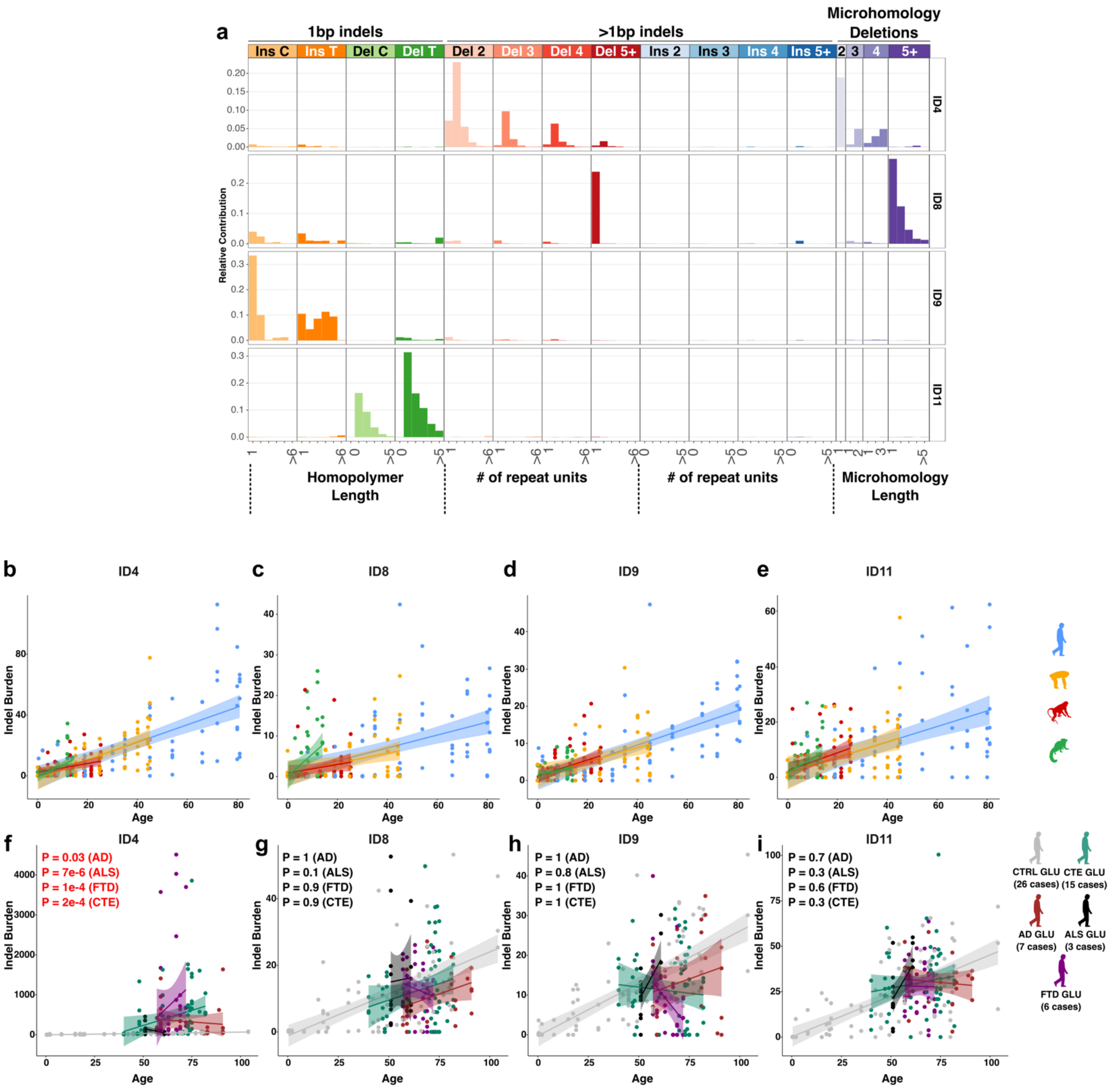


**Supplementary Figure 11: Indel mutational signatures across species and neurodegenerative diseases.** (a) COSMIC indel mutational signatures detected in this study's human dataset based on SigProfilerExtractor[60]. (b-e) Genomewide mutation burdens of indel signatures across species. (f-i) Genomewide mutation burdens of indel signatures across neurodegenerative signatures. P-values indicate significant excess in disease compared to control. CTRL: healthy control, CTE: chronic traumatic encephalopathy, AD: Alzheimer's disease, ALS: amyotrophic lateral sclerosis, FTD: frontotemporal dementia. Significant p-values (P<0.05) are shown in red.

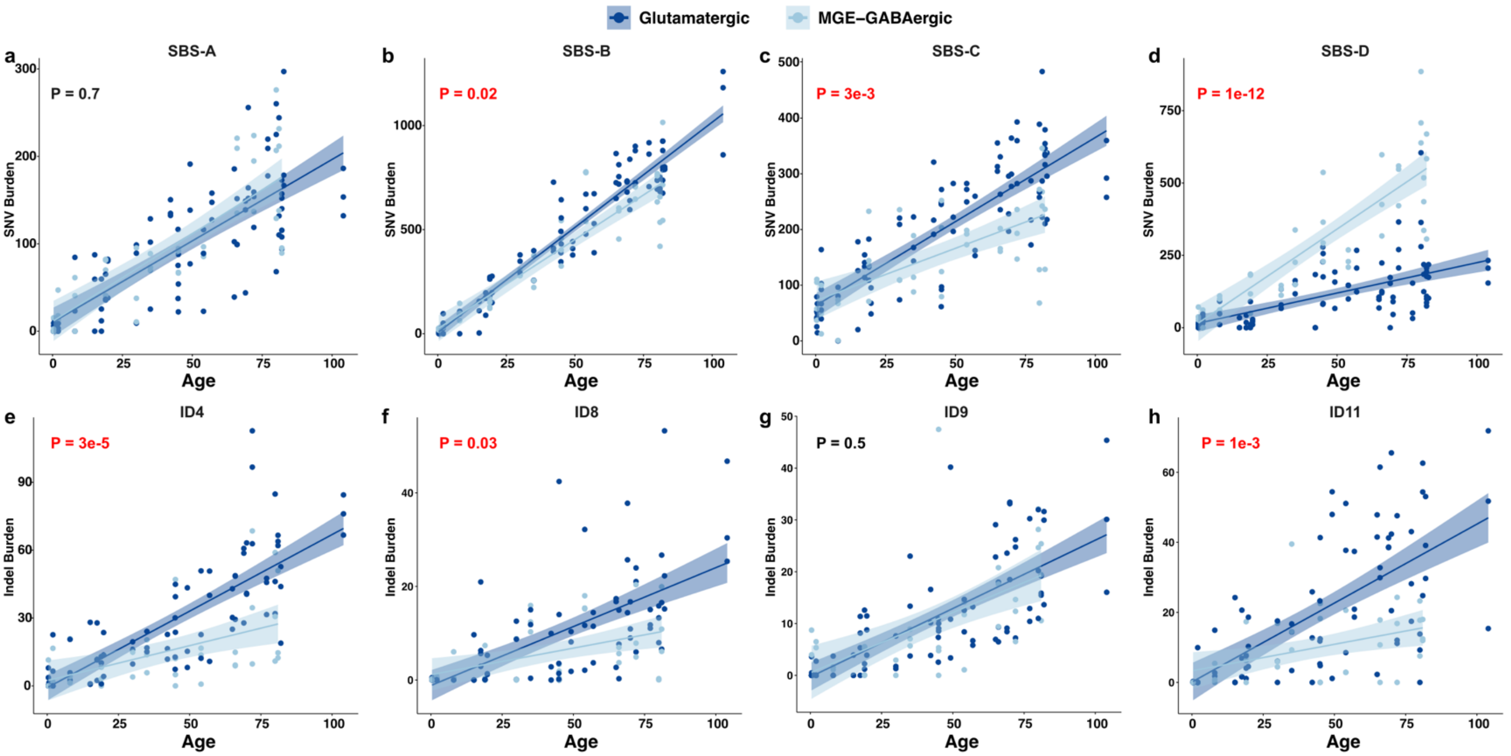


**Supplementary Figure 12: Cell type specific accumulation of mutational signatures.** (a-d) Genomewide mutation burden estimates per cell in SNV signatures and (e-h) indel signatures. Dark blue indicates glutamatergic, light blue indicates MGE-GABAergic cells. P values represent cell-type specificity of age-related change in mutation burden (burden ~ Age*CellType + (1|Subject)) for each mutational signature. P<0.05 are shown in red.

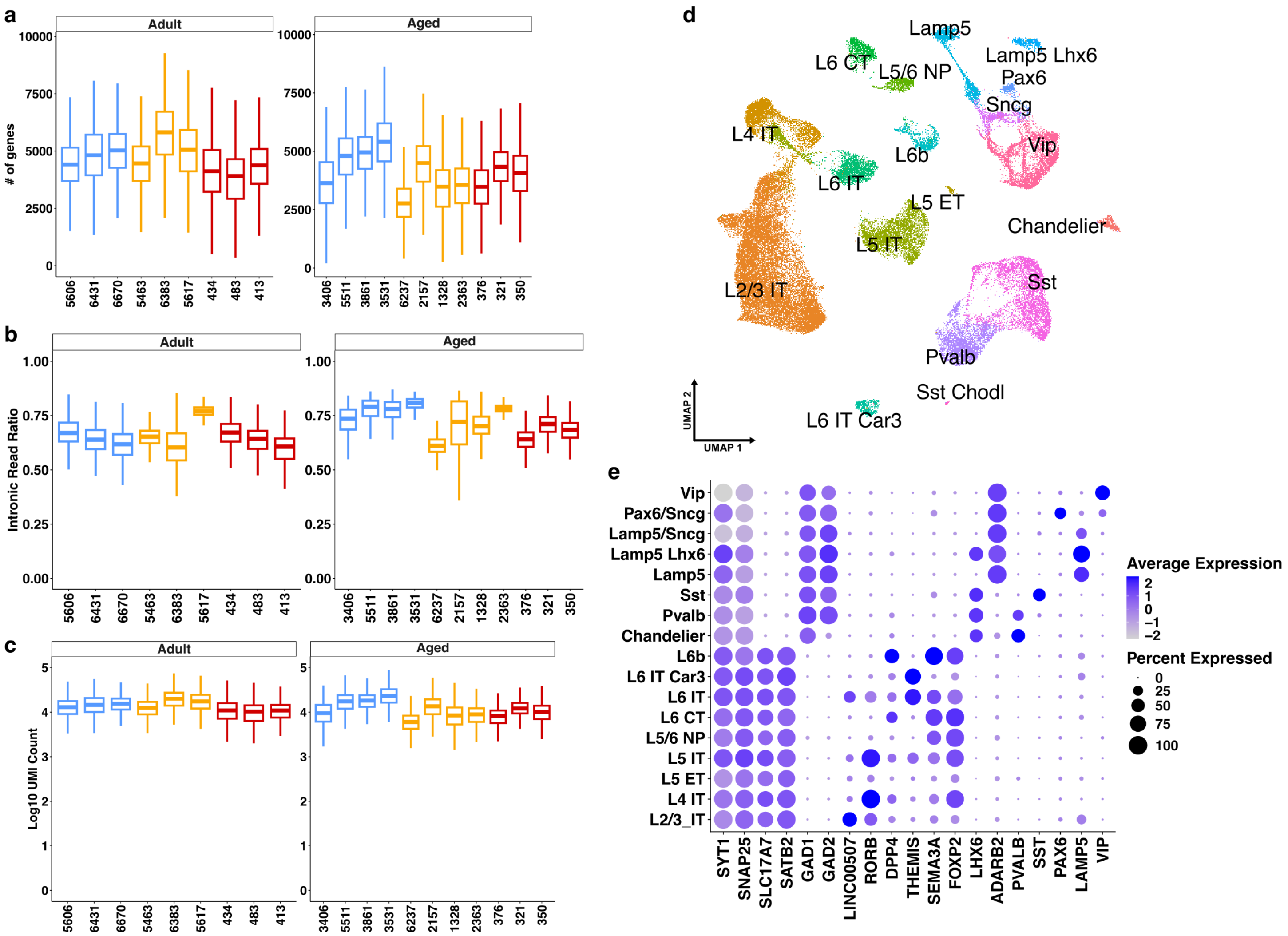


**Supplementary Figure 13: Additional plots on snRNA-seq across species and ages.** (a) Boxplot distribution of number of genes in all cells per sample per species. (b) Same as (b) for intronic read ratio (ratio of all reads that map to introns). (c) Boxplot distribution of $\log_{10}$ UMI count in all cells per sample per species. (d) Cell type annotations at neuronal subtype resolution displayed in the UMAP space. (e) Dot plots of transcript levels of marker genes (x-axis) per cell type (y-axis). Dot sizes represent percentage of cells that express each gene and colors represent normalized and z-transformed gene expression values across cell types.

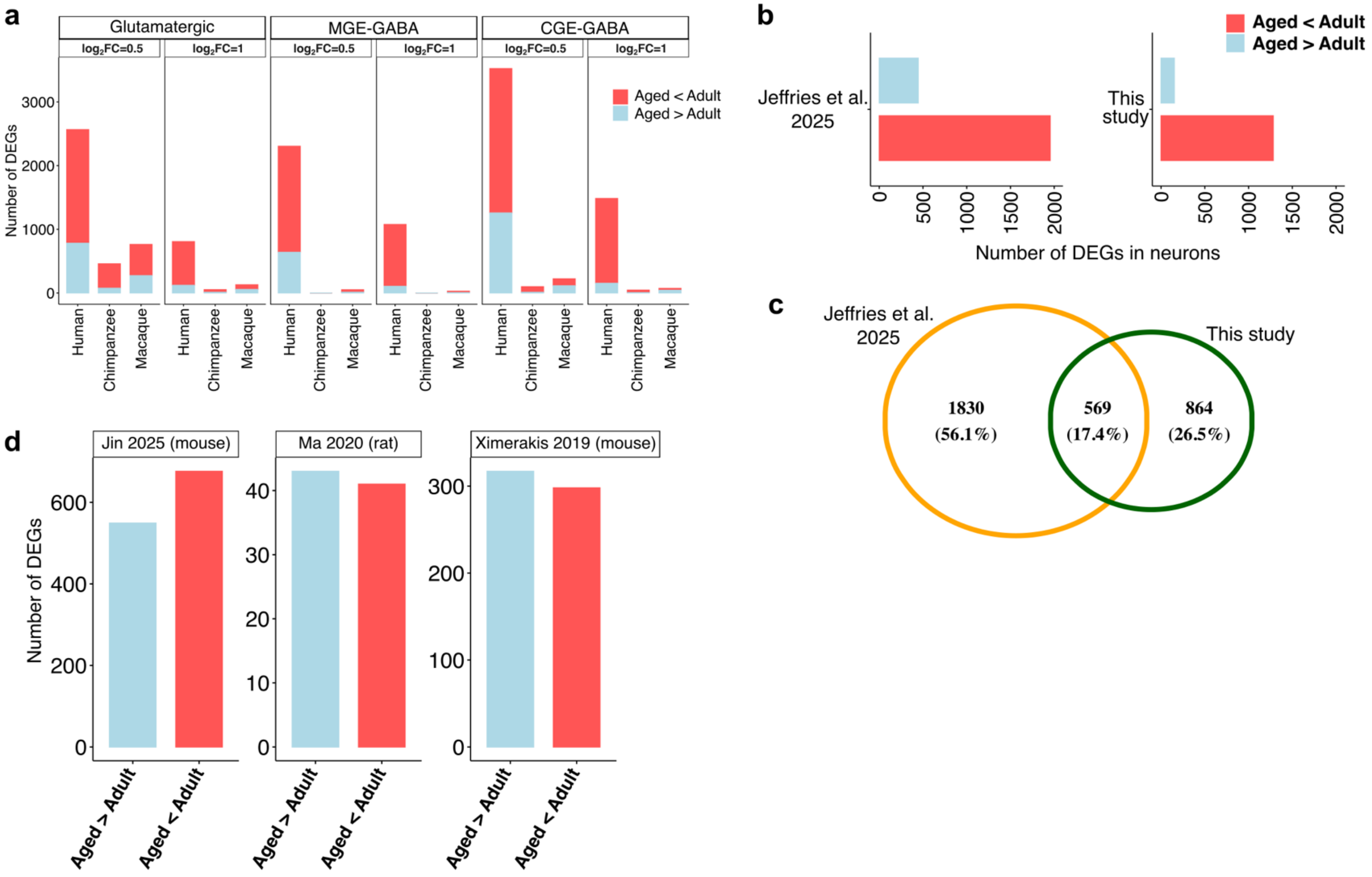


**Supplementary Figure 14: Additional analyses on age related DEGs.** (a) Number of DEGs between adult and aged individuals in human, chimpanzee rhesus macaque for each major cell type with the following cutoffs: absolute $log_2FC > 0.5$ and FDR < 0.05. (b) Number of DEGs between adult and aged groups in the neurons of an independent study from Jeffries et al.[29]. Right panel shows the pooled number of DEGs between adult and aged groups in this study for comparison. (c) Overlap of DEGs in Jeffries et al. and this study shown in panel b. An overlap considered if the change was in the same direction (e.g aged < adult in both datasets). (d) Number of DEGs between adult and aged groups in the neurons of independent studies performed on rodents. Number of DEGs were counted and plotted from the supplemental tables of each study for all external studies and no further modifications or analyses were performed.

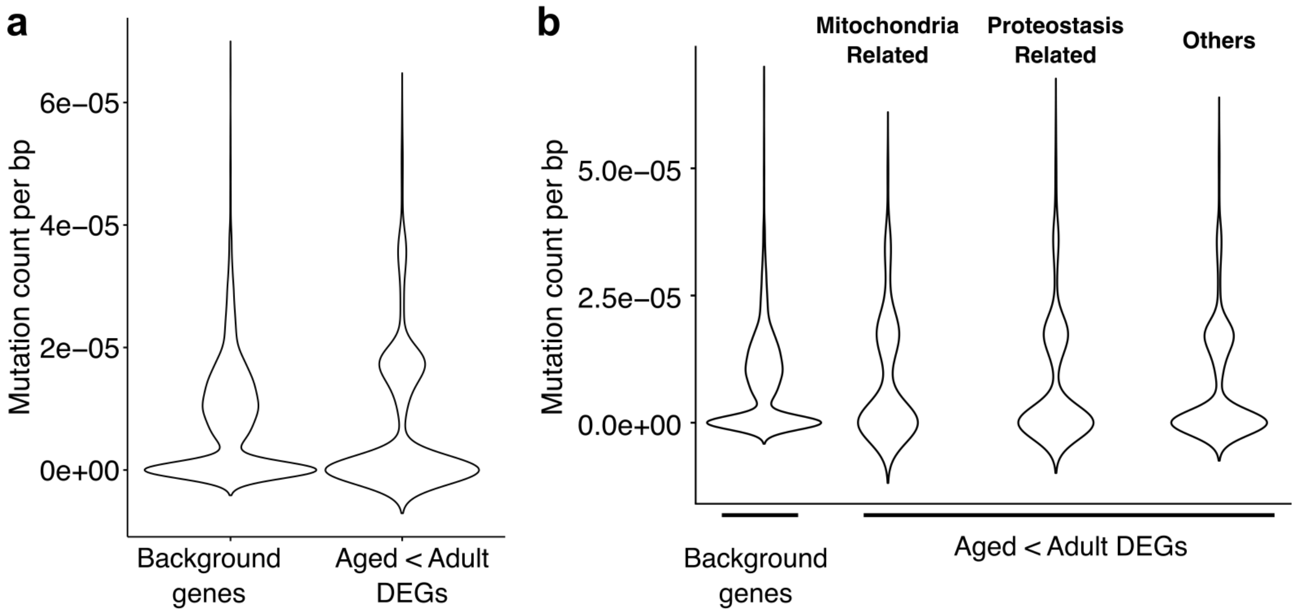


**Supplementary Figure 15: Normalized mutation counts for genes of interest and the background.** (a) Mutation (SNV or indel) count per bp for all DEGs classified as aged < adult or background. Background was all genes tested for differential gene expression but were not classified as a significant difference. (b) Same as panel a but for aged < adult DEGs classified as with mitochondria or proteostasis related functions and the rest (‘Others’).

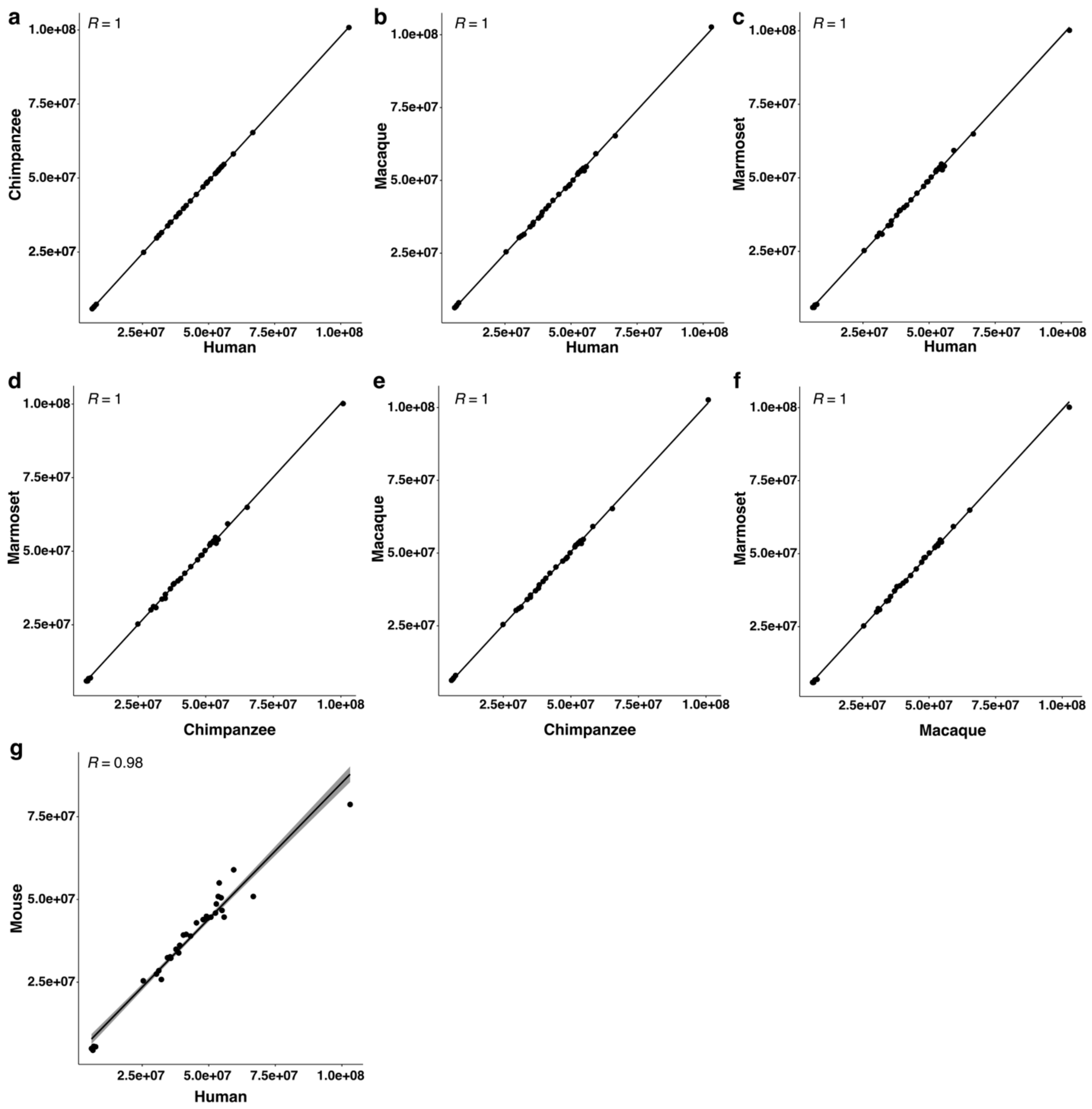


**Supplementary Figure 16: Comparisons of trinucleotide context counts across different species' genomes**. (a) Number of counts for each trinucleotide contexts are displayed on both axes for human and chimpanzee genomes. (b-g) Same as panel a for other species as shown in each plot. Pearson's correlation coefficient is shown on each plot with 'R'. Shaded area around the linear curve represents 95% confidence interval.

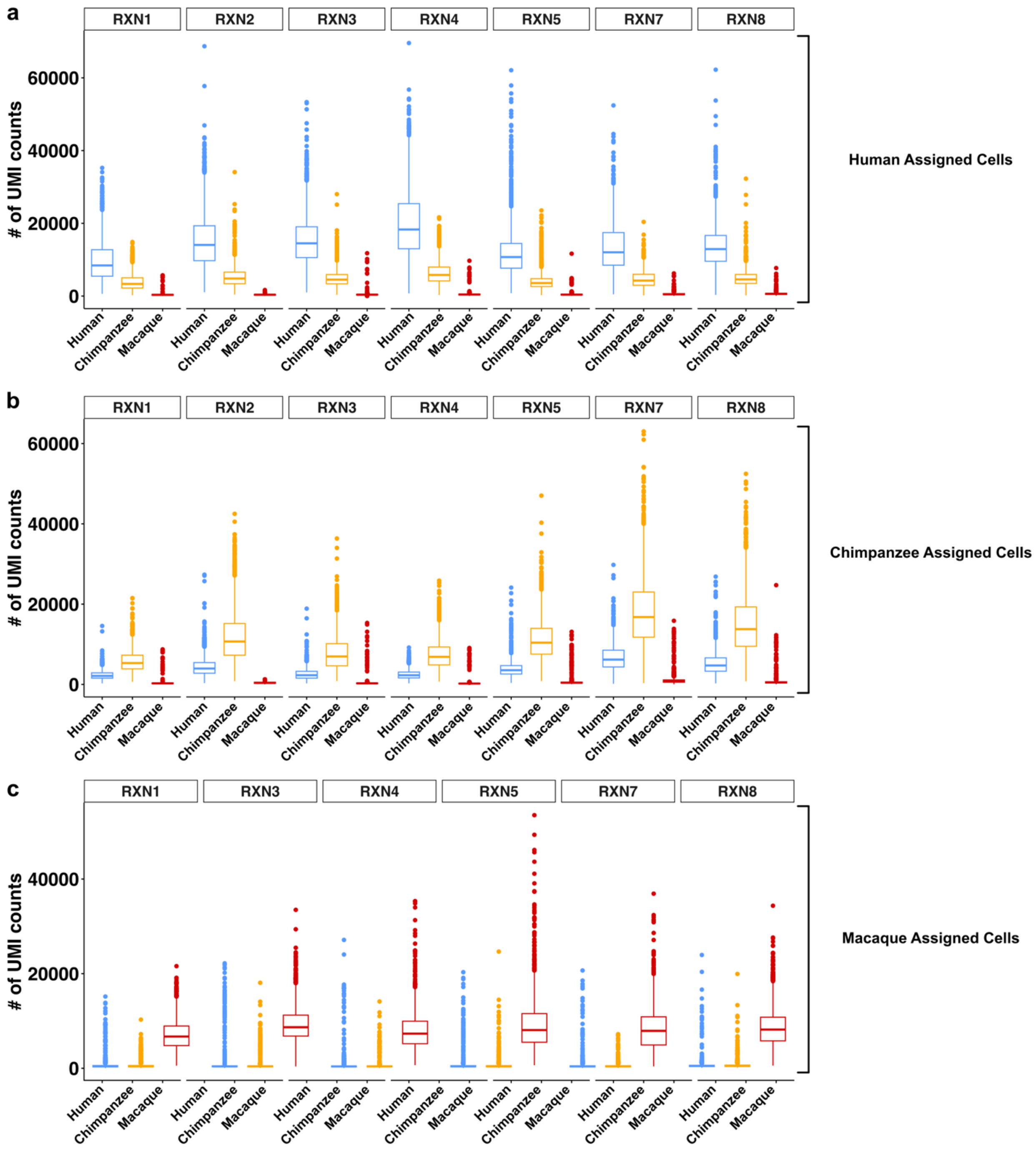


**Supplementary Figure 17: Comparing perfectly aligned UMI counts in different species.** Number of UMI counts for reads that have perfect (i.e without a mismatch) alignment to each species' own genome for cell barcodes that are assigned to (a) human, (b) chimpanzee or (c) rhesus macaque.
